# SARS-CoV-2 membrane protein biogenesis

**DOI:** 10.64898/2026.01.19.700281

**Authors:** Juan Ortiz-Mateu, Guy J. Pearson, Marina Rius-Salvador, Shiva Sedighian, Anna Pavlova, Josep Alonso-Romero, José M. Acosta-Cáceres, Ane Metola, María Jesús García-Murria, J. Mark Skehel, James C. Gumbart, Jeremy G. Carlton, Gunnar von Heijne, Manuel Mateo Sánchez-Del Pino, Luis Martínez-Gil, Ismael Mingarro

## Abstract

Viral protein biogenesis underpins every viral life cycle stage, and elucidating these processes could reveal fundamental principles of virus–host interaction, and vulnerabilities amenable to therapeutic targeting. Here we apply biophysical, molecular, and cell biology techniques to investigate the insertion, folding, and oligomerization of the SARS-CoV-2 M protein. We describe the sequential co-translational insertion of the hydrophobic core, and demonstrate that the cytosolic C-terminal domain undergoes slower adoption of its tertiary structure. Additionally, we characterize how the transmembrane domain bundle facilitates M-protein oligomerization. Our results reveal a hydrophobic residue cluster that is essential for protein folding and co-translational dimerization. Additionally, we identify the cellular machinery responsible for targeting and inserting the M protein into the ER membrane, and chaperones and cofactors that may contribute to proper folding.

## Introduction

Viral protein biogenesis underpins every stage of the viral life cycle. Elucidating these biosynthetic pathways can provide critical insights into virus–host molecular interactions and reveal exploitable nodes for antiviral development. Severe acute respiratory syndrome coronavirus 2 (SARS-CoV-2) is a positive-sense single-stranded RNA virus. Its genome encodes four structural proteins: the Spike (S), Envelope (E), Membrane (M), and Nucleo (N) proteins. Virion assembly is initiated within the ER–Golgi intermediate compartment (ERGIC), where the M protein serves as the central organizer, coordinating structural component recruitment,^1,2^ and driving membrane budding.^3^

The M protein is a dimeric type III membrane glycoprotein containing three transmembrane domains (TMDs) that connect with a cytosolic β-sheet domain.^2,4^ Structural analysis,^2,5,6^ have revealed short and long forms of the M protein. Transition from the long to the short form may facilitate virion assembly by inducing membrane curvature.^6^ Small-molecule inhibitors that block this transition have demonstrated strong antiviral activity,^7,8^ underscoring the therapeutic potential of targeting M-protein biogenesis.

Multipass membrane proteins are targeted to the ER when their signal peptide or first TMD is recognized by the signal recognition particle (SRP).^9–11^ Next, the SRP delivers the ribosome–nascent chain complex to the SRP receptor,^12,13^ allowing insertion, mainly by the Sec61 translocon.^14^

This insertion process is supported by accessory proteins. Among them, the translocon-associated protein (TRAP),^15^ aids in translocating secretory proteins and large luminal loops,^16–19^ the ER membrane complex (EMC), of the Oxa1 insertase family, accommodates unstructured N-terminal regions and inserts the first TMDs of proteins with short, non-, or negatively-charged luminal domains.^20–25^ Other accessory complexes, collectively termed the “multipass translocon” (MPT), play key roles in multipass protein biogenesis.^26–29^ The PAT, GEL, and BOS complexes function as chaperones and scaffolds, especially for low-hydrophobic TMDs.^30,31^ Interactions between EMC and BOS suggest that these components form a dynamic integrated holo-insertase system.^32^

Here we employ molecular and cell biology approaches to dissect the ER-targeting, membrane integration, folding dynamics, and oligomerization behavior of SARS-CoV-2 M protein. Force profile analysis and mammalian topology assays reveal fast sequential TMD insertion and slower cytosolic folding. We described a co-translational dimerization mediated by the TMD bundle and confirm that a cluster of hydrophobic residues is essential for protein dimerization and function. We further demonstrate that TMD2 is the key structural domain supporting this oligomerization. Finally, we identify the cellular machinery involved in targeting and inserting the M protein into the ER membrane, and chaperones and cofactors that may contribute to proper folding. These findings define the molecular principles governing M-protein biogenesis and reveal potential host-dependent checkpoints that could be leveraged to disrupt coronavirus assembly.

## Results

### Co-translational folding of M protein

To investigate the co-translational insertion and folding of the SARS-CoV-2 M protein, we employed in-depth force profile analysis (FPA), which enables residue-level resolution of membrane integration events during translation (See **Methods** section).^33–36^ This is achieved using a protein arrest peptide (AP) that binds in the ribosome exit tunnel, thereby stalling translation.^37^ If the nascent chain is subjected to a strong pulling force (PF) the AP is removed and translation proceeds at its normal rate.^38^ Therefore, the AP can be used as a “molecular force sensor” to probe co-translational force-generating processes. **Figure 1A** shows that a TMD inside the ribosome exit tunnel does not generate PF on the nascent chain, but generates a strong force during membrane insertion.^33^

**Figure 1.**
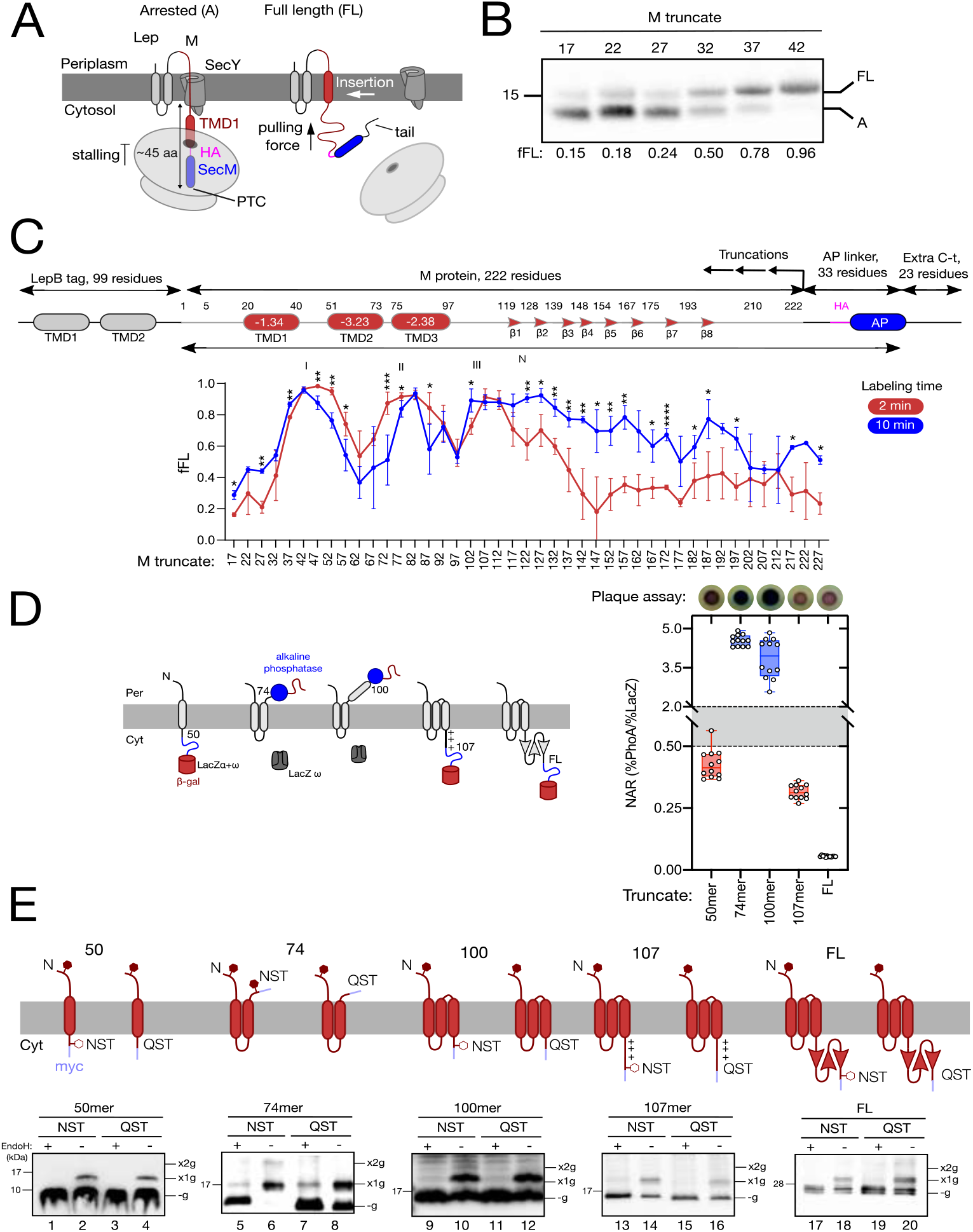
Co-translational folding of SARS-CoV-2 membrane protein. (A) Schematic of force profile analysis (FPA). Translation of a truncated protein (red), fused to an arrested peptide (AP, blue) and C-terminal extension (black), induces ribosomal stalling when the C-terminal residue of the AP occupies the peptidyl transferase center (PTC) without membrane insertion. When a sufficiently hydrophobic TMD reaches the membrane insertion site, ~45 residues from the PTC, the generated PF relieves the stall. (B) Representative FPA result. SDS-PAGE analysis of immunoprecipitated, [^35^S]-Met-labeled constructs (identified by truncated lengths). PF is quantified as the fraction of full-length protein (f_FL_), calculated from the ratio of intensities of the full-length (FL) and arrested (A) products. The fFL values reflect the PF generated by TMD1 membrane insertion. (C)(Top) Schematic representation of force profile constructs. M protein was fused to a 4-residue SGSG linker, 12-residue HA tag, 17-residue SecM AP, and 23-residue tail. TMD regions (red) and β-strand-rich domain (red arrows) are indicated. Each predicted TMD’s hydrophobicity is indicated as the ΔG value for insertion, calculated with ΔG Predictor. N denotes the length between M’s first residue and the arrest protein (AP)’s last residue. (Bottom) FPs for M protein with 2 min (red) or 10 min (blue) radioactive labeling. Above each data point: *p* values for unpaired t-tests. Individual variance was assumed for each condition. \**p* > 0.05, \*\**p* > 0.01, \*\*\**p* > 0.001, \*\*\*\**p* > 0.0001. (D) (Left) Truncated M-protein insertion in E. coli cells. Truncated M protein was fused to the pho-lac dual reporter to assess membrane topology. Cytosolic topology (Cin) allows β-galactosidase α and ω fragment reconstitution in the cytoplasm, yielding detectable enzymatic activity. Periplasmic topology (Cout) allows proper folding and activity of the alkaline phosphatase moiety. (Right) Experimental determination of M-truncate membrane topology. (Top) E. coli DH5α cells expressing different M/Pho-lac fusions were plated on indicator medium with Red-Gal (for β-galactosidase activity) and X-Pho (for phosphatase activity). Blue colonies indicate high phosphatase activity. (Bottom) Quantitative enzymatic assays of selected M/pho-lac fusions. Box plot shows normalized activities ratios (NAR): blue boxes for phosphatase activity; red boxes for β-galactosidase activity. NAR values of >2 indicate periplasmic location, 2–0.5 transmembrane location, and <0.5 cytosolic localization. Solid dots represent individual replicates. (E) (Top) Truncated M protein was fused to a C-terminal glycosylation site. Constructs were expressed in HEK293T cells with a non-natural glycosylation-competent site (NST) or a control site that cannot be glycosylated (QST) plus a C-terminal c-Myc tag (blue). Cout topology allows NST-site glycosylation (red hexagons), whereas Cin topology does not (white hexagons). (Bottom) M-truncate expression in HEK293T cells reveals different glycosylation states. Cell lysates were treated with EndoH (+, odd lanes) or untreated (−, even lanes) before western blot analysis. Bands corresponding with non-glycosylated (−glyc), singly glycosylated (+1x), and doubly glycosylated (+2xglyc) forms are indicated.

For the FPA, M protein was truncated from its C-terminus at 5-residue resolution. Each construct was fused to the SecM AP, followed by a linker and a C-terminal tail derived from *E. coli* LepB (**Figure 1A**). We also fused LepB N-terminal region upstream of the M protein, to increase expression and ensure efficient targeting to the SecYEG translocon.^34^ Constructs were expressed in *E. coli*, followed by a short pulse with [^35^S]-Met. M truncates were immunoprecipitated and analyzed by SDS-PAGE (**Figure 1B**). Quantitation of the bands corresponding to the arrested and full-length forms of each construct allowed calculation of the fraction of FL protein f_FL_ = I_FL_/(I_FL_+I_A_), where I_FL_ and I_A_ are the respective band intensities.

**Figure 1C** (bottom; red curve) shows the full FPA. Three distinct peaks are observed (I, II, and III), each with maximum fFL values at truncation positions ~47, 82, and 107. These peaks support a model of sequential co-translational insertion of all TMDs, exhibiting peak amplitudes near 1, indicating strong PF during translation. This force correlates with the TMDs’ high hydrophobicity, which likely facilitates their membrane insertion.^39,40^ Peaks I and II occur when the N-terminal end of each TMD is approximately 60 residues from the peptidyl transferase center, consistent with other membrane proteins.^33,35^ In contrast, the maximum fFL for peak III was observed when the N-terminal end was about 65 residues away, suggesting that residues down-stream of TMD3 influence its insertion.

The loops between TMD1 and TMD2, and between TMD2 and TMD3, exhibit moderately elevated fFL values. Additionally, the region following TMD3, including the initial segment of the β-strand-rich cytoplasmic domain, generates measurable PF (M-truncations 122–137). To investigate these phenomena, we recorded a second FPA with increased labeling from 2 to 10 min (**Figure 1B**, blue curve). Giving the AP more time to be withdrawn from its binding site usually yields increased fFL values for regions with low hydrophobicity or other constraints.

The fFL values associated with the loop between TMD1 and TMD2 were significantly reduced by a 10-min pulse. This is possibly because 2 minutes are insufficient for the topological rearrangements of TMD2, thus recording higher fFLs as the TMD is adopting its N_in_-C_out_ topology. Extending the labeling time may allow these rearrangements, thereby diminishing the PF exerted by the nascent chain at 10 min. The fFL values remained elevated for the region between TMD2 and TMD3, suggesting a folding or dimerization event involving TMD1 and TMD2. Increasing the labeling time augmented the fFL values in the β-strand-rich cytoplasmic region, demonstrating slow folding kinetics for this domain.

Using a 10-minute labeling time shifted the maximum fFL associated with peaks I and III from truncates 47 to 42, and 107 to 102, respectively. This indicates that, given sufficient time, TMD1 and TMD3 could fully partition into the lipid bilayer earlier, without requiring residues down-stream of the TMD. Sequence inspection reveals arginine residues at position 44, following TMD1, and at positions 101, 105, and 107, shortly after TMD3. The “positive-inside rule”^41^ states that positively charged residues are typically retained on the cytoplasmic side of the membrane; thus, these arginine residues may facilitate TMD1 and TMD3 insertion, with each adopting an Nt^out^/Ct^cyt^ orientation.

To validate the M-protein folding dynamics described by the FPA, we sought to prove the sequential TMD insertion using a bacterial *pho-lac* dual-reporter assay (See **Methods** section)^42–44^ (**Figure 1D**). We engineered a series of chimeric constructs with the dual reporter fused after each TMD (residues 50, 74, and 107), and after the full-length protein. Additionally, we generated a construct with the reporter fused at residue 100, excluding R101, R105, and R107. Cells expressing constructs fused at residue 50 yielded red colonies, indicating LacZα activity and cytoplasmic orientation. Fusion at residue 74 yielded blue colonies, consistent with alkaline phosphatase activity and periplasmic orientation (**Figure 1C**). When the reporter was located at the C-terminal end of the full-length M protein, cytoplasmic orientation was observed (**Figure 1D**). Fusion after the third TMD (residue 100) produced blue colonies whereas fusion at residue 107 yielded red colonies (**Figure 1C**). These results confirm that positively charged residues facilitate TMD3 insertion.

To investigate sequential insertion of the TMDs of M protein in mammalian cells, we designed a series of truncated constructs, each harboring a C-terminal glycosylation reporter immediately after each TMDs. The reporter comprised either a glycosylation competent (Asn-Ser-Thr, NST) or incompetent (Gln-Ser-Thr, QST) sequence, enabling discrimination between cytosolic and luminal orientations based on glycosylation^45,46^ (**Figure 1E**). All constructs retained the native N-terminal glycosylation site (N^5^GT) (Figure S1A).

All constructs containing only TMD1 (50mer) showed one faint higher-molecular-weight band. Endo H treatment confirmed the glycosylation origin of this band, verifying that N-terminal is in the ER lumen, while the C-terminal remains cytoplasmic (**Figure 1E**). Truncates 100mer and 107mer exhibited a singly glycosylated form, confirming successful TMD3 membrane insertion. These findings suggest that in mammalian cells, the intrinsic hydrophobicity of TMD3, possibly aided by chaperones or membrane insertion machinery, is sufficient to drive proper membrane integration.

In the glycosylation assay, the full-length protein showed a single higher-molecular-weight band (**Figure 1E**, lanes 10 and 14). In contrast, the 74mer truncate was both singly and doubly glycosylated. The unmodified form of the 74mer was nearly undetectable, suggesting reduced glycosylation efficiency at the native N-terminal site, compared to the engineered C-terminal site (~95% versus ~55%). Incorporating the Omicron-specific D3G and Q19E mutations into the N-terminus did not affect glycosylation efficiency (**Figures S1A** and **S1C**). This implies that distinct glycosylation mechanisms may govern modification at each site (**Figure 1E**, lane 6).^47^

### Characterizing M-protein co-translational oligomerization via TMD interactions

Previous studies show that the M protein forms functional dimers within viral particles.^5^ The transmembrane dimer interface comprises two interacting bundles, where TMD1 from one monomer aligns between TMD2 and TMD3 of the opposing monomer.^2,4^ To explore the M protein’s self-association properties, we first utilized the BLaTM assay, a tool designed to asses TMD-TMD interactions.^48^

While we detected did not homotypic or heterotypic interactions between TMD1 and TMD3 (**Figures 2B** and **2D**), TMD2 showed strong specific interactions with both TMD1 and TMD3 (**Figure 2C** and **2D**). Also, no TMD interacted with the negative controls (Figure S2).

**Figure 2.**
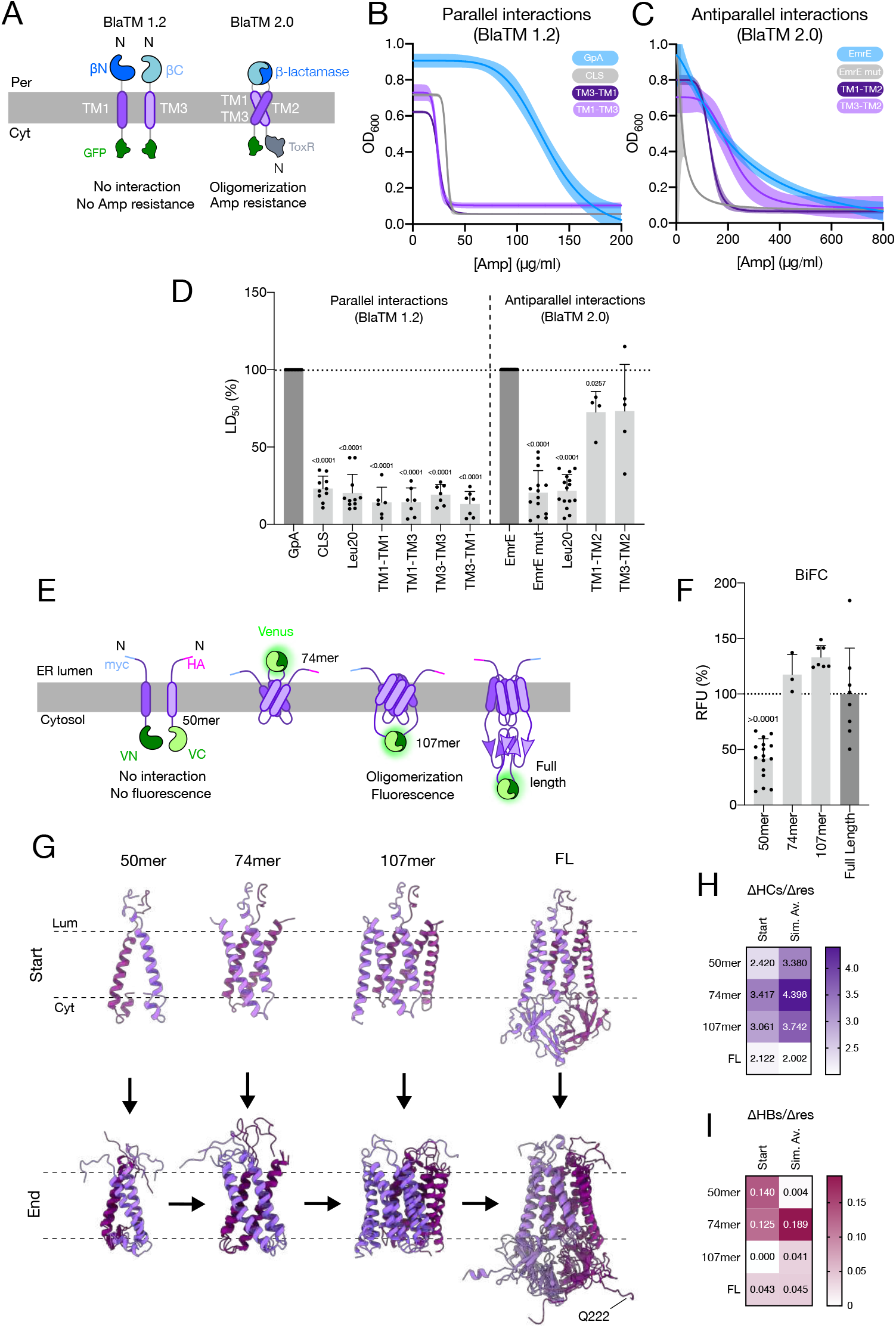
Experimental model for M-protein co-translational oligomerization. (A) To investigate TMD interactions within the SARS-CoV-2 M protein, we employed the BlaTM assay. Productive TMD–TMD interactions reconstitute the β-lactamase enzyme, enabling bacterial growth on selective media. The BlaTM 1.2 system was used to evaluate parallel homo- and heterotypic interactions between TMD1 and TMD3. The BlaTM 2.0 system enables testing of antiparallel heterotypic interactions between TMD2 and TMD1 or TMD3. (B) In the BLaTM assay, the LD_50_ of the antibiotic ampicillin indicates the strength of the TMD–TMD interaction. Representative examples of dose– response curves used to calculate LD_50_ values. Positive control: GpA TMD homodimer (blue). Negative control: CLS TMD (gray). (C) Same as in (B) but for the BlaTM 2.0 assay. Positive control: TMD homodimer of EmrE (blue). Negative control: a mutant that disrupts this interaction (EmrE mut) (gray). (D) LD_50_ values for M TMD–TMD interactions. Values are normalized against the positive controls (dark gray; set to 100%). Data are presented as mean ± SD. Solid dots represent individual experiments (*N* = 5–15). Above bars: *p* values below the significance threshold (< 0.05), calculated using one-sample t-tests against a reference value of 100. (E) M-protein oligomerization was assessed via bimolecular fluorescent complementation (BiFC) assay. The N-terminal and C-terminal ends of the VFP (VN and VC, respectively) were fused to the C-termini of M-protein truncates. The specific residues in each truncate are indicated. (F) BiFC relative fluorescence units (RFU) for M truncates and full-length homo-interactions. All values are normalized to the mean RFU of the full-length M-protein interaction (dark gray; set to 100%). Mean ± SD is shown. Solid dots represent individual experiments (*N* = 3–15). Above bars: *p* values below the significance threshold (<0.05) calculated using one-sample t-tests against a reference value of “100”. (G) Structures of M-protein-dimer constructs from our simulations, from left to right: 50mer, 74mer, 107mer, and full-length. Top representations correspond to the start point of the simulations. Bottom representations show an overlay of three independent replicates’ end-points. (H) Quantification of hydrophobic contacts (HCs) from MD simulations. Results presented as the ratio of the average increase in hydrophobic contacts (ΔHCs) per residue increment (Δres) across the 400-ns simulation (Sim. Av.) and at the initial timepoint (Start) for each simulated dimer. Numerical values are displayed on the heatmap. HCs calculated at the simulation start point are also shown. (I) Same as in (H) for hydrogen bonds (HBs).

We also used BiFC in eukaryotic cells to investigate M-protein oligomerization (See **Methods** section) (**Figure 2E**).^49–51^ Co-expression of VN- and VC-tagged full-length M proteins yielded strong fluorescence, confirming that the system reported M-protein oligomerization (**Figure 2F**). The 50mer construct failed to produce fluorescence (**Figure 2F**), despite confirmed protein expression (**Figure S3**). The 74mer (containing TMD1 and TMD2) and 107mer (containing TMD1–3) successfully reconstituted VFP fluorescence, implying that the hydrophobic core, particularly TMD2, is sufficient to drive oligomerization (**Figure 2F**).

Additionally, we assessed the hetero-oligomerization potential among truncated variants with compatible membrane topologies. Only the 107mer and full-length constructs reconstituted VFP fluorescence (**Figure S4**), highlighting the importance of the complete TMD bundle for M-protein dimerization.

To further investigate the TMD bundle’s role in M-protein dimerization, we employed AlphaFold multimer^52^ to generate structural models of potential dimers for the truncated and full-length M proteins. We performed molecular dynamics (MD) simulations on the dimeric full-length and truncated M proteins, to assess their structural stability and interaction dynamics (**Figure 2G**). The 50mer dimers exhibited the fewest hydrophobic contacts and hydrogen bonds per added residue (**Figures 2H** and **2I**), suggesting weaker interactions. Full-length M dimers also showed a decreased hydrophobic contact per added residue ratio, likely because the soluble C-terminal domain (CTD) did not significantly contribute to membrane-embedded interactions.

Despite dynamic differences, the 107mer and full-length dimers remained closely associated throughout the simulations (**Figure 2G**). In contrast, a pronounced partial separation was observed around residue 40 in the 50mer, which was reduced in the 74mer, supporting that TMD2 plays a critical role in dimer formation. In concordance with previous experimental results, even further stabilization was observed by the complete TMD bundle (**Figures S5A** and **B**).

Analyzing the root mean square fluctuation of atomic positions revealed peak near residue 40 across all dimers (**Figures S5A** and **S5B**). Closer inspection identified N41 and R42 as the primary contributors. These hydrophilic residues point outward toward the membrane head groups, resulting in increased flexibility compared to the rest of the structure. Moreover, our data suggest that flexibility is amplified upon partial dimer strand separation.

### A hydrophobic cluster at the transmembrane interface stabilizes M-protein dimers

We next sought to identify the specific residues critical for dimer stabilization. MD simulations of the full-length M protein revealed a cluster of hydrophobic residues at the dimer interface, which remained in close contact (**Figures 3A** and **3B**). This cluster includes L22 and F26 in TMD1 of one monomer, interacting with F65 in TMD2 and M84 in TMD3 of the opposing monomer. These four residues are at the monomer–monomer interface, with L22 and F65 buried within the TMD bundle, and F26 and M84 exposed to the lipid environment, enabling protein–lipid interactions. This cluster was observed across all available M-protein cryo-EM structures,^2,4,6^ including its short and long conformations, with inter-residue distances ranging from 3.4–4.6 Å (**Figures 3A** and **S6**).

**Figure 3.**
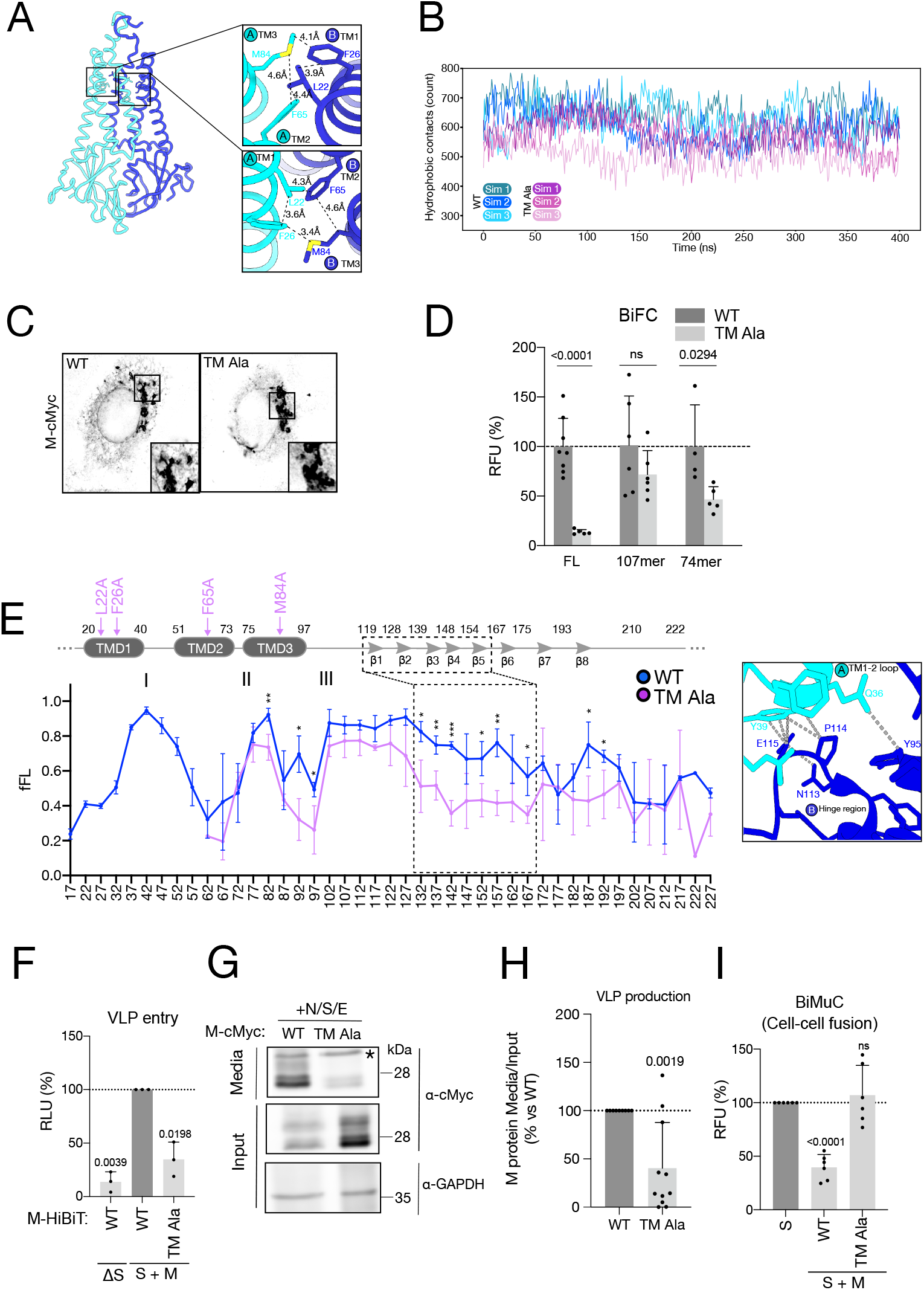
Functional characterization of a transmembrane hydrophobic cluster governing M dimerization. (A) (Left) AlphaFold prediction of M-protein structure, highlighting a cluster of hydrophobic residues involved in dimeric interface stabilization. Distances between residues are indicated in Å. (B) Hydrophobic contacts over time between the two chains of the full-length M dimer in MD simulations. Each line corresponds to independent simulations. Blue indicates wild-type M-protein simulations; purple indicates M-TMD-Ala mutant simulations. (C) Subcellular localization of the M wild-type and M-TMD-Ala mutant. Representative images of VeroE6 cells transfected with plasmids encoding myc-tagged M wild-type and M-TMD-Ala mutant. (D) BiFC assay showing oligomerization levels of wild-type and TMD-Ala M-protein constructs. Relative fluorescence units (RFU) are shown, normalized to the mean RFU of the respective wild-type versions (dark gray; set to 100%). Data presented as mean ± SD. Solid dots represent individual experiments (*N* = 5–9). Above each comparison: *p* values below the significance threshold (<0.05), calculated using pairwise t-tests against a reference value of 100. (E) FP of the wild-type (blue) and TMD-Ala (purple) mutant versions of M protein. Both profiles correspond to a 10-min labeling time (as in **Figure 1B** for wild-type). Above data points: *p* values for unpaired t-tests. Individual variance was assumed for each condition. \**p* > 0.05, \*\**p* > 0.01, \*\*\**p* > 0.001, \*\*\*\**p* > 0.0001. Above the fFL data: schematic representation of M secondary structure. Dotted-line square highlights the region of a significant difference in fFL, corresponding to contacts between the hinge region and TMD1–TMD2 loop. (F) BiMuC assay showing fusogenic protein activity with or without wild-type M and TMD-Ala mutant. Values are normalized against the positive control (dark gray; set to 100%). Data are presented as mean ± SD. Solid dots represent individual experiments (*N* = 5). Above each bar: *p* values below the significance threshold (<0.05), calculated using one-sample t-tests against a reference value of 100. (G) Expression and VLP incorporation of codon-optimized wild-type M and TMD-Ala mutant, N, S, and E proteins in HEK293T cells. HEK293T cells were transfected with plasmids encoding codon-optimized versions of wild-type M protein and the TMD-Ala (cMyc tagged), along with N, S, and E proteins. Cell lysates (Input) and media containing VLPs were analyzed by immunoblotting with antisera against the c-Myc tag. GAPDH was used as a loading control and marker for the cellular fraction. Asterisk (*) indicates a non-specific band. (H) Western blot quantification of M protein. Band intensities from cell lysate (Input) and media (VLP-containing fraction) were quantified, and the input-to-media ratio was calculated. All values are normalized to wild-type M (dark gray; set to 100%). Data are presented as mean ± SD. Solid dots represent individual experiments (*N* = 11). Above each bar: *p* values below the significance threshold (<0.05), calculated using one-sample t-tests against a reference value of 100.

To assess the contributions of these residues to M protein dimerization, each was substituted with alanine (M TMD-Ala), and MD simulations were performed. M TMD-Ala showed a reduced average hydrophobic contacts (552 ± 36) compared to wild-type (629 ± 19) (**Figure 3B**), indicating that this hydrophobic cluster plays an important role in dimer stabilization.

To test the relevance of the identified hydrophobic interface, we generated an M-protein expression plasmid carrying the aforementioned alanine substitutions. The TMD-Ala variant successfully reached the ERGIC compartment, similar to wild-type (**Figure 3C**). However, M-TMD-Ala showed impaired oligomerization (**Figure 3D**). When these mutations were introduced into the 74mer and 107mer truncations, oligomerization was still impacted, although less than in the full-length mutant (**Figure 3D**).

To further probe the effects of the mutations we recorded an FP of the M-TMD-Ala protein. The amplitudes of peaks II and III were reduced for the mutant, despite the higher hydrophobicity of TMD2 and TMD3 (**Figure 3E**). Furthermore, the fFL values following TMD2 were markedly reduced in the M-TMD-Ala, suggesting that TMD2 insertion enables rapid engagement in inter-molecular interactions. We also observed decreased fFL values in the 102–122 truncations, supporting previous findings that TMD3 contributes to the oligomerization of the TMD bundle. A reduction is also observed in 137–157 truncations, reflecting influence of the TMD bundle in CTD dimerization (**Figure 3D**). These results support the hypothesis that C-terminal β-strand domain dimerization depends on prior TMD bundle dimerization. The alanine substitutions might disrupt the dimerization interface, inducing conformational change of the TMD bundle, potentially leading to steric clashes between the soluble CTDs, and contributing to the impaired oligomerization of the full-length protein.

To assess the functional consequences of hydrophobic cluster mutations, we first employed HiBiT-tagged virus-like particles^53^ (VLPs; see **Methods**). Briefly, VLPs carrying HiBiT-tagged M protein (wild type or TMD-Ala mutant), along with the S protein, were used to infect cells expressing LgBiT luciferase (**Figure 3F**). Successful infection reconstitutes NanoLuc, resulting in luminescence.^54^ Elevated luminescence was observed with the wild-type M protein, whereas VLPs containing the TMD-Ala mutant produced only background signal, suggesting that virion formation or viral entry is impaired in the mutant.

Additionally, we investigated whether the M-TMD-Ala mutant could support assembly into virus-like particles (VLPs, **Figures 3G** and **3H**).^55–57^ We co-expressed wild-type or M-TMD-Ala with the other structural proteins in HEK293T cells. Despite comparable expression levels (**Figure 3G**), the M-TMD-Ala mutant resulted in markedly reduced VLP production (**Figure 3H**), highlighting this hydrophobic cluster’s critical role in M-mediated assembly and budding.

Finally, we employed a cell–cell fusion assay (BiMuC, **Figure 3I**),^58,59^ which allowed evaluation of the M protein’s role in recruiting the S protein to the ERGIC (See **Methods** section).^1,60^ As reported M co-expression retained S in the ERGIC, thereby inhibiting membrane fusion.^1,60^ Coexpression of the M-TMD-Ala fully restored fusion activity to levels comparable to those with S protein alone (**Figure 3I**), indicating that the mutated hydrophobic residues in M-TMD-Ala, and likely the M-protein dimerization, are essential for M–S interaction and S protein retention in the ERGIC.

### Vicinal proteome of truncated and full-length M protein variants

To identify cellular factors involved in M-protein insertion and folding in the ER, we investigated the local environment during its biogenesis. We employed a proximity labeling strategy, fusing an HA-TurboID tag to the full-length protein and truncated M variants, to obtain snapshots of the biogenesis process (**Figure 4** and **S7A**) (See **Methods** section).

**Figure 4.**
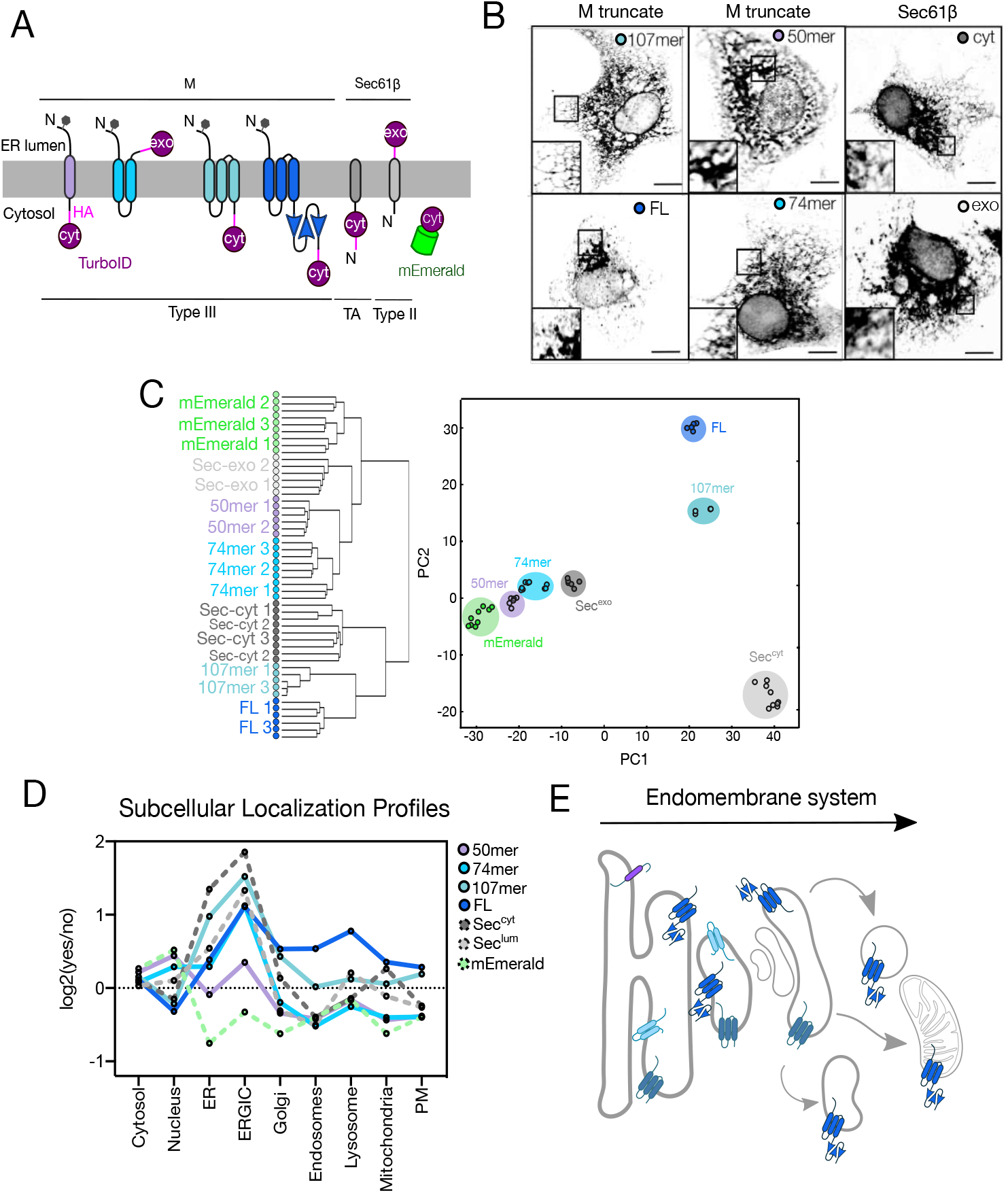
A proximity labeling approach to study the M protein’s vicinal proteome. (A) Construct design and topology. Type III substrates comprising truncated M variants were fused to a C-terminal HA-TurboID tag. Constructs with Cout topology positioned TurboID in the ER lumen (exo), while those with C_in_ topology retained TurboID in the cytosol (cyt). Controls included Sec61β fused to TurboID at the N-terminus (cytosolic orientation; tail-anchored substrate) or C-terminus (luminal orientation; type II substrate), and mEmerald-TurboID as a cytosolic reference. (B) Subcellular construct localization. VeroE6 cells transfected with plasmids encoding M or Sec61β-HA-TurboID were fixed and stained with anti-HA antibody. Images are representative of over 10 analyzed images per condition. (C) Clustering and dimensionality reduction. (Left) Hierarchical clustering of proteomic samples based on label-free quantification (LFQ) values, plotted using average Euclidean distance. (Right) Principal component analysis (PCA) on median-averaged data across three experimental replicates per sample. (D) Subcellular enrichment analysis. Profiles display the log_2_ ratio of average abundance of proteins assigned to specific subcellular compartments (based on UniProt annotations) relative to all other proteins. Ratios above zero indicate enrichment within the indicated compartment. (E) Schematic representation of the primary subcellular localization of full-length M protein and its truncated variants (50mer, 74mer, and 107mer), based on proximity labeling and enrichment analysis.

The cytosolic mEmerald fluorescent protein served as a non-membrane control (**Figure 4A**). We also used tail-anchored Sec61β subunit of the ER translocon, fused to TurboID at its N-terminus (cytosolic orientation, Sec^cyt^) or C-terminus (luminal orientation, Sec^exo^) (**Figure 4A**).

The controls allowed to differentiate true partners from background signals. Comparing the 74mer construct with Sec^exo^ provided insight regarding luminal partners specific to M’s membrane-embedded regions (**Figure 4A**). All constructs localized to intracellular membranes, except the mEmerald control (**Figure 4B**). The Sec^cyt^ and Sec^exo^ reference proteins displayed strong ER localization. Full-length M exhibited a Golgi-like distribution, whereas the truncated variants’ pattern was consistent with ER localization (**Figure 4B**).

We performed a TurboID-mediated biotinylation reaction and isolated the biotinylated proteins. Efficient biotinylated protein enrichment, and their successful release following on-bead digestion was confirmed (**Figures S7B** and **S7C**).

Identified proteins were filtered to select *bona fide* tagged hits (**Figure S7D**). A cluster analysis of the hits reveals the 107mer variant clustered closely with full-length M, whereas the 50mer and 74mer grouped together (**Figure 4C**). Among control samples, Sec^exo^ and mEmerald clustered near the 50mer and 74mer, while Sec^cyt^ was separated from all other samples.

To further evaluate the cellular context of the biotinylated proteomes, we performed initial functional categorization based on UniProt’s “subcellular compartment” annotations (**Figures 4D** and **4E**). The 50mer sample showed low specific membrane compartment association, reflecting a mislocalized distribution. TMD2 introduction increased association with ER and ERGIC-resident proteins. The 107mer displayed strong labeling of ER and ERGIC proteins, but reduced association with proteins of downstream compartments. The full-length M protein was enriched for ERGIC-associated proteins and showed significant labeling of proteins in later secretory pathway stages, including the Golgi apparatus, endosomes, lysosomes, mitochondria, and plasma membrane (**Figures 4D** and **4E**). Among the controls, both Sec constructs primarily labeled ER and ERGIC proteins, while mEmerald showed low overall labeling of membrane-associated proteins. These findings align with our confocal imaging results (**Figure 4B**), demonstrating that the M CTD is necessary for its anterograde transport through the secretory pathway. This role of the CTD had been characterized previously for other coronaviruses such as MERS-CoV M protein,^61^ but not in SARS-CoV where the trafficking signal had been suggested to be at the TMDs.^62,63^

Next, we investigated the proximity of the full-length M and truncates to the ER-localized membrane protein targeting and biogenesis machinery (**Figure 5A**). We detected both subunits of the signal recognition particle receptor (SR) (**Figures 5B, S8A**, and **S8B**). SRα showed strong labeling across all samples, likely due to its high expression. Nonetheless, the lowest intensity was identified with the Emerald construct. TMD1 constitutes the signal anchor that might target M to the ER via SRP:SR complex formation, exhibiting an intensity of 16-fold higher than the median in the shorter, 50mer sample (**Figure S8A**). SRβ displayed a pattern similar to SRα across all samples, albeit at lower intensities (**Figure S8B**).

**Figure 5.**
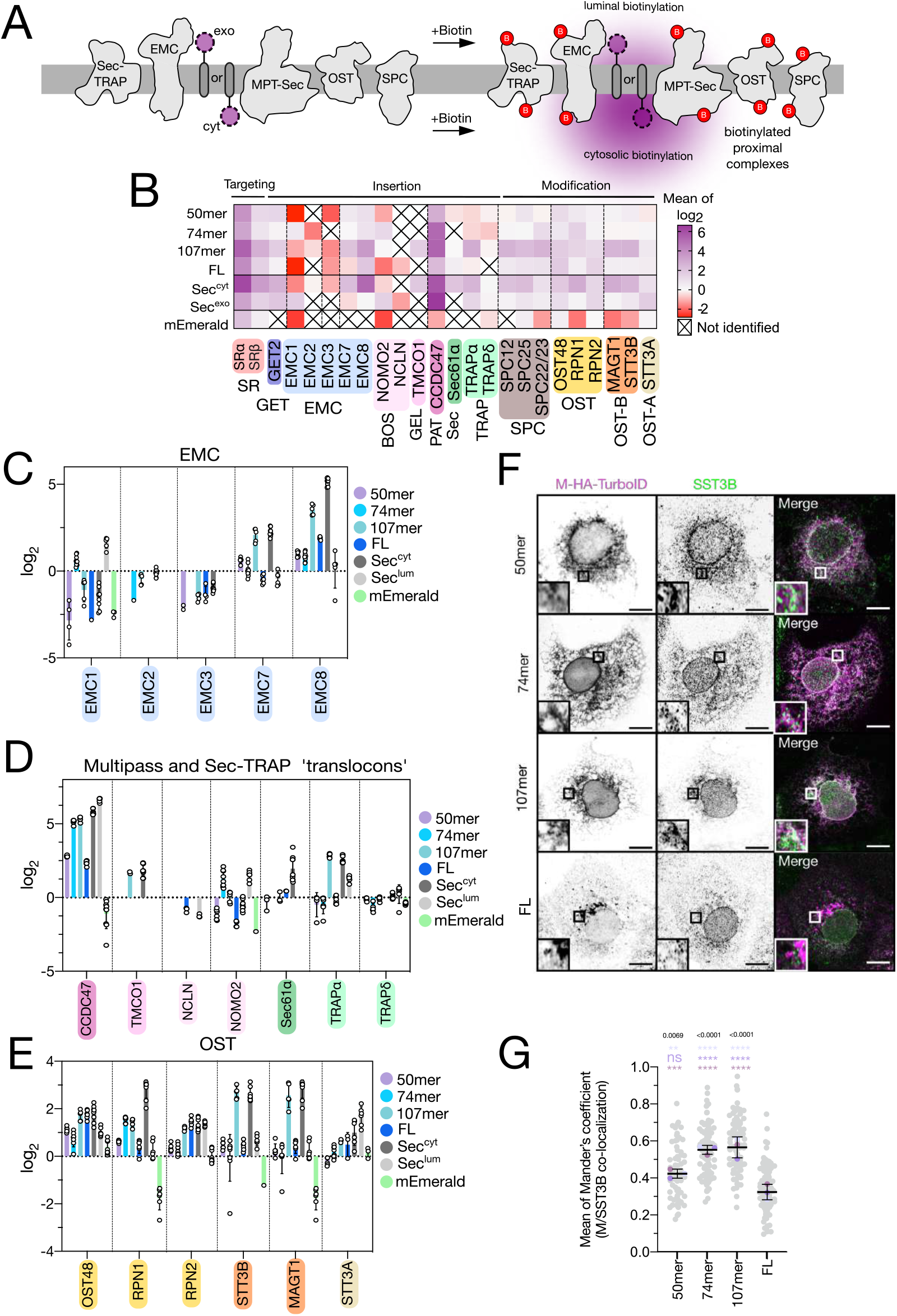
Screening of proximity-labeled human insertases assisting M biogenesis. (A) Upon biotin addition for 20 minutes, the *exo* and *cyt* substrates biotinylate the luminal or cytosolic domains of membrane protein complexes, respectively. (B) Heatmap representation of abundance distribution values of biogenesis factors near the substrates tested in the proximity labeling approach. Gradient represents the mean value of the median-normalized abundance in log_2_ scale. (C) Bar plots show detailed abundance of the different EMC subunits detected in our MS data. Protein abundance is represented as median normalized in log_2_ scale. Mean ± SD is shown. Solid dots represent an *N* of 4–9. Dotted line indicates the median value. (D) Same as in (C) for components of the multipass and TRAP-Sec translocons. (E) Same as in (C) for components of the human OST complexes. (F) Representative images of VeroE6 cells transfected with plasmids encoding truncated versions of M-HA-TurboID, fixed and stained with antisera raised against HA tag and SST3B. Squared boxes indicate colocalized M and SST3B. Images representative of ≥70 captured images in each case. (G) Mander’s coefficient calculation for quantifying cells displaying SST3B co-localization of newly synthesized M-HA-TurboID proteins imaged in (F). *N* = 70, 95, 94, and 124 imaged cells (grey dots), respectively. Three independent experiments with means highlighted in overlay (purple, pink, and blue dots). Mean ± SD displayed over the individual dots. Significance determined by unpaired t-tests against the full-length sample.

Next, we assessed the proximity labeling data for ER insertase complexes (**Figures 5B, 6C**, and **6D**).^32^ These insertases and accessory complexes also stabilize early-inserted TMDs, while subsequent ones emerge from the ribosome. Out of the known human insertase complex subunits^14,27,28,31,32,64,65^ 13 were detected in our dataset (**Figure 5B** and **Table S1**). Only four were weakly detected in the mEmerald control, confirming membrane localization of the other baits. GET2, a tail-anchored insertase subunit, was most enriched in the tail-anchored Sec^cyt^ control (6-fold over median), and showed above-median enrichment in Sec^exo^, 50mer, and 107mer samples (**Figures 5B** and **S8C**).

Among insertases, the EMC complex is the primary candidate for mediating insertion of the M protein’s TMD1, which adopts an N^exo^ orientation and contains a short negatively charged luminal loop and high TMD hydrophobicity. The luminal subunit EMC1 was highly enriched in the C^exo^ samples (Sec^exo^ and 74mer) (**Figures 5B, 6C**, and **S8D**). The EMC subunits most frequently enriched across samples were EMC7 and EMC8, a cytosolic subunit involved in client recognition^23^ (**Figures 5B, 6C, S8E**, and **S8F**). EMC8 was strongly biotinylated by 107mer, full-length M, and the Sec^cyt^ constructs (4- to 32-fold over median), in agreement with their membrane topology, and showing reduced labeling in 50mer and 74mer variants.

Interactions between the EMC and BOS complexes^32^ suggest that MPT members, such as PAT, GEL, and/or BOS, may assist in TMD2 and TMD3 insertion after EMC-mediated TMD1 insertion. Our data support this hypothesis. Within the GEL complex, TMCO1-which may participate in insertion of multipass substrates as highly enriched in 107mer and Sec^cyt^ samples (4-fold over median) (**Figures 5B, 6D**, and **S8H**).

Regarding the PAT complex, CCDC47 was among the most abundantly labeled proteins overall, showing changes of 4- to 128-fold over the median across samples (**Figures 5B, 6D**, and **S8I**). Its labeling pattern widely varied, with strong enrichment seen in 74mer, 107mer, and both Sec61β constructs, but lower labeling in 50mer and fulllength M (an 8- to 32-fold difference between groups). This pattern suggests that CCDC47 may preferentially assist with TMD2 and TMD3 insertion,.^27,28,66^ Asterix, which facilitates low-hydrophobicity TMD insertion, was not detected, aligning with the highly hydrophobic nature of M protein’s TMDs.

The BOS complex, with an unknown role in the MPT, was less abundantly labeled. The luminal BOS subunit NOMO2 was labeled in the 74mer (4-fold over median) (**Figures 5B, 6D**, and **S8G**), in agreement with its membrane topology. NCLN was detected at low levels, and TMEM147 was not detected.

Finally, we assessed Sec61 translocon components. Sec61α was enriched in the Sec^cyt^ sample (4-fold over median), but not in the luminal Sec^exo^ variant, and it appeared at or near median levels in M protein samples (a 4-fold change) (**Figures 5B, 6D**, and **S8J**). TRAPα, mainly a luminal subunit, was highly enriched in 107mer and Sec^cyt^ (8-fold over median), and moderately labeled in Sec^exo^ (2-fold) (**Figures 5B, 6D, S8K**). TRAPδ was detected at lower intensity, and the remaining translocon subunits (Sec61β, γ, TRAPβ, and TRAPγ) were not detected. This indicates that the Sec61 channel is not as proximal as other Oxa-1 family insertases from the M multipass truncated variants, while it is proximal to the Sec^cyt^ control.

We next focused on proteins that may act as molecular “modifiers” of the M protein (**Figure 5B**). Our first target was the signal peptidase complex (SPC). Since the TMD1 functioning as a signal anchor, rather than a cleavable signal peptide, we did not expect significant proximity to SPC components. However, our data revealed strong enrichment of the integral adaptor subunits, SPC12, SPC25, and SPC22/23 (**Figures S9A, S9B**, and **S9C**), while the catalytic subunits SEC11A and SEC11C were not detected.This pattern suggests a possible structural or scaffolding interaction, rather than enzymatic processing.

Recent findings indicate that M protein undergoes polylactosamine glycosylation, post-translationally catalyzed by the OST-B complex.^47^ We identified six OST subunits with distinct labeling profiles (**Figures 5B** and **6E**). Among the shared components of OST-A and OST-B,^67^ we detected OST48 (**Figure S9D**), Ribophorin I (RPN1, **Figure S9E**), and Ribophorin II (**Figure S9F**, RPN2)

We detected robust enrichment of the OST-B-specific catalytic subunit STT3B and its adaptor, the redox-active chaperone MAGT1. Both showed significantly higher abundance in 107mer and Sec^cyt^ samples, up to 8-fold over median (**Figures 5B, 6E, S9G**, and **S9H**). These results were validated by immunostaining and co-localization analyses, which confirmed the proximity of endogenous STT3B to the 74mer and 107mer constructs, but not the shortest 50mer truncate (**Figures 5F** and **6G**).

STT3A, the catalytic subunit specific to OST-A, was not significantly enriched in the M protein samples. Its labeling was only slightly increased in the luminal Sec^exo^ control, consistent with its association with Sec61 (**Figures 5E** and **S9I**). Furthermore, its associated adaptor proteins, DC2 and KCP2, were entirely absent from the biotinylation dataset. These findings strongly support that SARS-CoV-2 M protein glycosylation may be mediated by the OST-B complex.

### A subset of ER chaperones is proximal to the M-protein TMD bundle

We next examined whether ER-resident chaperones might be involved in assisting M-protein folding following TMD insertion. We generated a network based on Pearson correlation coefficients between the log_2_-fold enrichment profiles across Turbo-ID baits, retaining only positive correlations above 0.8 (**Figures 6A** and **S10**). Clustering analysis of this network revealed 30 distinct clusters of at least five proteins each. One prominent sub-network comprising three clusters, with similar average abundance patterns, exhibited notable enrichment in the Gene Ontology Biological Process category “protein folding in the endoplasmic reticulum” (**Figures 6A, 6B** and **Table S2**).

**Figure 6.**
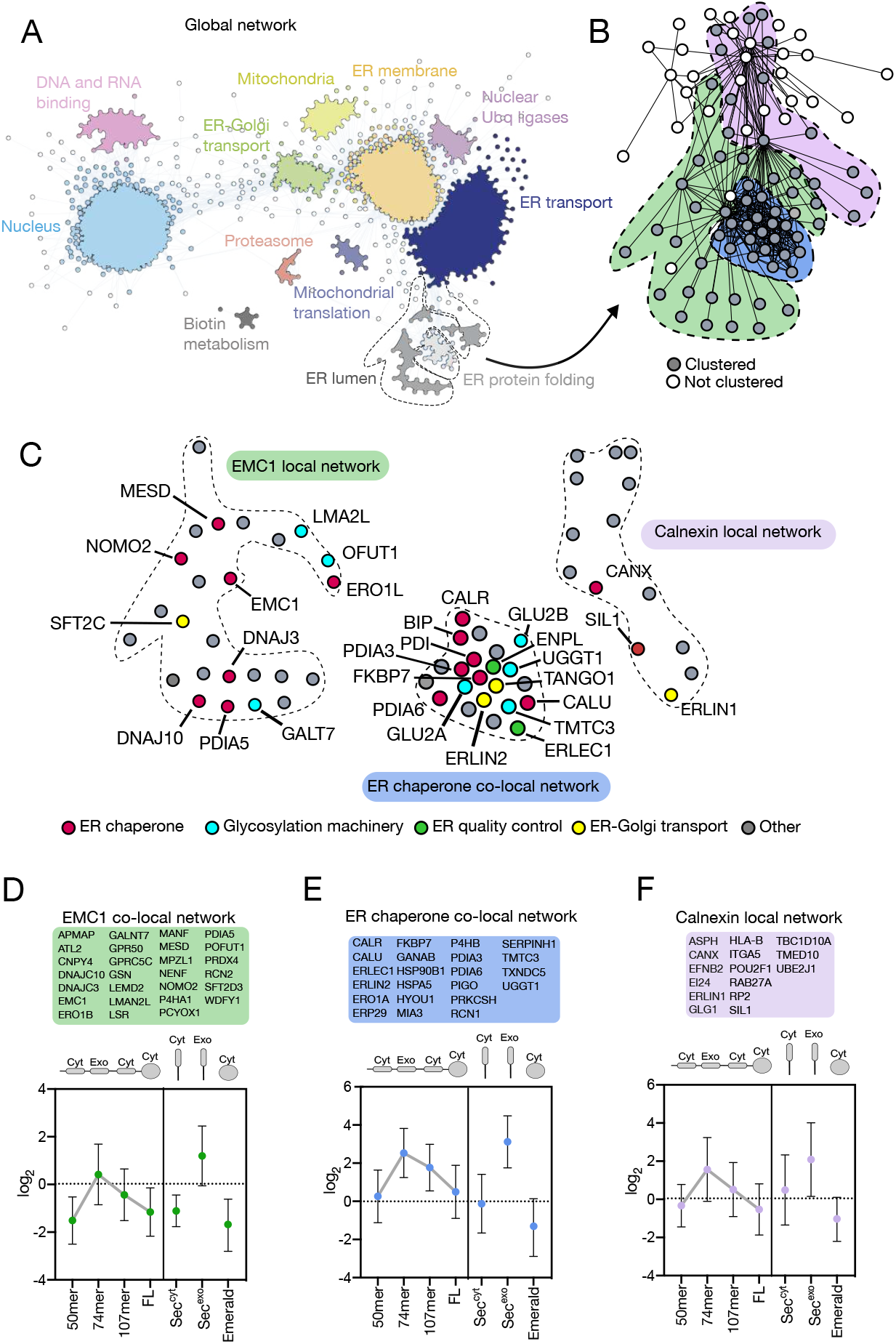
Identification of proximal co-factors mediating M stability. (A) Protein correlation network. Pearson coefficients among proteins were calculated, and positive correlations with coefficients of >0.8 were used to generate the network. The main protein clusters are colored, and the labels indicate their main biological process or cellular locations. (B) Details of the endoplasmic reticulum protein folding sub-network. MCL algorithm identified three clusters that are shaded with different colors. White nodes are not part of the three clusters. (C) Details of proteins of interest found within the three clusters. Colors highlight different functions relevant for ER protein biogenesis and homeostasis. (D) (Top) List of proteins clustered within the EMC1 local network. (Bottom) Abundance profile showing the average cluster protein abundance. Mean ± SD; N = 3 biological replicates, up to 9 technical replicates. (E) Same as in (C) but for the ER chaperone co-local network. (F) Same as in (C) but for the Calnexin local network.

Detailed functional enrichment analysis confirmed that all three clusters contain ER chaperones^68^ (**Figure 6C**). The central cluster was designated “ER chaperone co-local network”, includes key folding enzymes, such as calreticulin, protein disulfide isomerases, and calumenin (**Figure 6E**). This cluster also comprises glycoprotein-processing enzymes involved in N-glycosylation, trafficking factors and ER quality control components. In the cluster named “Calnexin local network”, the central node is calnexin (CANX), and ERLIN1 is the only other associated quality control component. The cluster named “EMC1 local network” also features chaperones and co-factors, (**Figure 6F**). This cluster also contains glycosylation-related proteins, ER-Golgi trafficking mediators involved in glycoprotein cargo transport and MPT insertase subunits.

All three clusters were most strongly labeled by Sec^exo^, consistent with their primarily luminal localization and roles in ER protein quality control. Similarly, the 74mer truncate, which places TurboID in the ER lumen, showed robust labeling of the proteins in all three clusters. Although these chaperones have more limited cytosolic tails for biotinylation, the 107mer truncate still displayed stronger proximity labeling for the ER chaperone co-local network, than the cytosolic Sec^cyt^ control, indicating specific enrichment of ER chaperones around multipass transmembrane substrates (**Figure 6D**).

## Discussion

### A model for SARS-CoV-2 M-protein biogenesis

Despite significant advances, many aspects of membrane protein biogenesis remain poorly understood,^69,70^ particularly relative to viral systems. Understanding these mechanisms is critical, as biogenesis errors can impair viral infectivity, and specific steps represent potential targets for antiviral intervention. Here we provide a detailed characterization of the co-translational insertion, folding, and assembly of the SARS-CoV-2 M protein, offering key insights into its synthesis and maturation (**Figure 7**).

**Figure 7.**
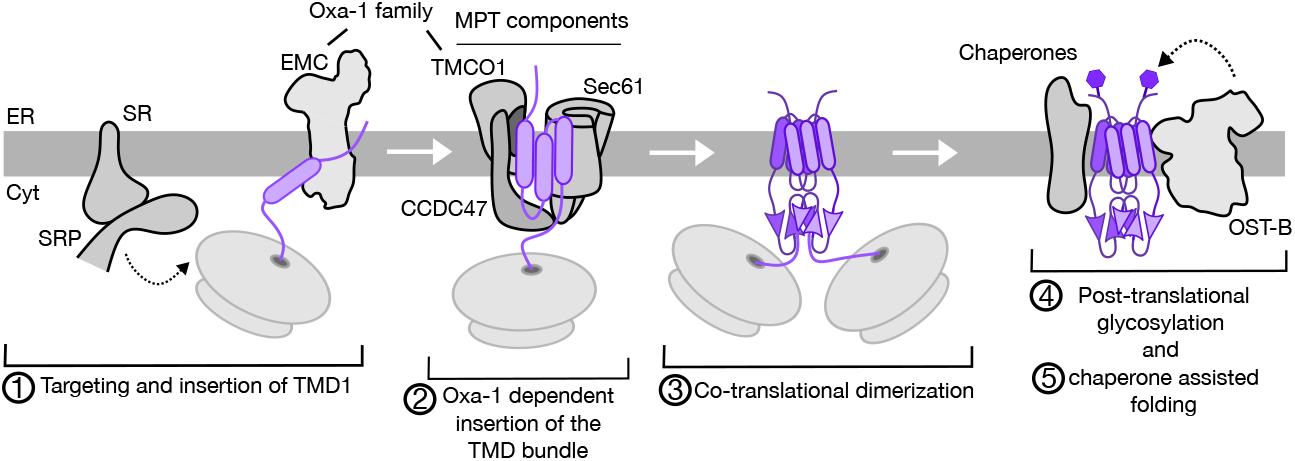
A model of SARS-CoV-2 M-protein biogenesis. M protein is targeted to the ER membrane by the SRP, following hand-off by this complex upon binding its receptor (SR). TMD1 acts as a signal anchor, inserted by an Oxa-1 family insertase, presumably the EMC complex (1). Next, the ribosome docks into Sec61 channel, and the rest of the TMD bundle is inserted by the side of the channel, with aid of other chaperones of the MPT (CCDC47 and TMCO1) (2). Upon TMD bundle insertion, it dimerizes while the cytosolic domain is translated in the cytosol (3). M dimers are post-translationally glycosylated by the OST-B complex (4), and then folded with help from different ER chaperones, like calnexin (5).

### Targeting and insertion of SARS-CoV-2 membrane protein

Different potential biogenesis pathways could explain the insertion of a multipass membrane protein without large luminal loops but highly hydrophobic TMDs: (1) TMD1 insertion by an Oxa1-superfamily insertase (presumably EMC), followed by hand-off of TMD2 and TMD3 to the MPT; (2) TMD1 insertion via Oxa1, with subsequent hand-off to the Sec61 translocon; or (3) insertion of the entire TMD bundle by either an Oxa1 insertase or the Sec61 translocon.

#### Targeting and insertion of TMD1

The biophysical characteristics of M’s N-terminal region support the model of Oxa1-mediated TMD1 insertion after hand-over from the SRP–SR complex.^24^ However, the prominent hydrophobicity of all three TMDs could hinder their compatibility with the partially hydrophilic vestibules of the Oxa1 members and MPT co-factors, like EMC3 in the case of EMC and TMCO1, or Asterix in the case of the MPT. Our data support Oxa1-dependent TMD1 insertion.

*In vitro* assays have shown that EMC can insert short unstructured N-terminal domains handed off from the SRP before their translocation into the ER lumen.^24^ The shortest truncate tested in this study can insert in the ER, adopting a N_exo_–C_cyt_ topology (**Figures 2D, 4B, 4D**, and **4E**). Consistently, in the proximity labeling assay, we detected the SRP receptor, and Oxa-1 insertase EMC (subunits 7 and 8) as proximal partners of the 50mer (**Figures 5B, S8A, S8B, S8E**, and **S8F**).

However, in terms of protein stability, the 50mer truncate is poorly glycosylated (**Figure 1E**), fails to oligomerize (**Figures 3E, 3G, 3H**, and **S4**), and does not stably localize in the ER (**Figures 4B, 4D**, and **4E**). Instead, it appears to accumulate near the nuclear envelope, and shows proximity to ER quality control system components (**Figure S10**). This indicates that, although the 50mer truncate can be targeted and inserted by EMC, its correct localization depends on the rest of the protein.

#### Oxa-1-dependent TMD bundle insertion

The next step in M biogenesis relies on insertion of TMD2 and TMD3. The two peaks in the FPA relating to TMD2 and TMD3 insertion (**Figure 1C**) indicates that TMD2 insertion is independent of TMD3, although both are almost contiguous in the sequence. Proper membrane insertion and topology were confirmed for 74mer and 107mer truncates in bacterial (**Figure 1D**) and human cells (**Figure 1E**). These truncates localized to the ER (**Figure 4B**) and biotinylated ER-resident MPs (**Figures 4D** and **4E**). We detected several EMC subunits with high LFQ intensities (**Figures 5B** and **S9**),. EMC2—known to facilitate TMD hand-off to the EMC vestibule via interaction with EMC8/9—is proximal to the 107mer (**Figures 4B** and **4C**). The holdase chaperone EMC8^23^ showed strong proximity labeling in our experiments, especially with the 107mer (**Figure 5C**), supporting its role in stabilizing the M-protein TMD bundle during insertion.

Our proximity labeling data showed biotinylation of several MPT-associated components. The BOS subunit NOMO2 is proximal to 74mer and to Sec^exo^. CCDC47 is labeled across all constructs, with the highest proximity to 74mer and 107mer truncates containing up to TMD2 and TMD3, respectively. The Oxa1 homolog TMCO1 (a GEL subunit) is detected in 107mer and in control Sec^cyt^ samples (**Figure S9H**). In contrast, Sec61α is detected mainly in Sec^cyt^, while Sec61β and γ are absent (Figure S9J), suggesting limited engagement between M protein and the full Sec61 complex.

A genome-wide CRISPRi screen^32^ revealed just two insertase-related genes affecting M stability: Sec61α and EMC2. The lack of effect from Sec61β and Sec61γ suggests that Sec61α’s involvement may be limited to ribosome docking at the ER membrane, rather than active TMD insertion. Moreover, this could indicate that functional compensation by different insertases.

We propose that M protein might versatilely adapt to different Oxa1 insertases and require the EMC complex and/or the GEL complex, with assistance of the PAT subunit CCDC47, which might be blocking the translocon channel to redirect the hydrophobic nascent chain to these alternative machineries.

### SARS-CoV-2 M-protein dimerization

Structural studies using cryo-electron microscopy have revealed that M proteins dimerize via domain-swapped TMDs and cytoplasmic domains, forming a “mushroom-shaped” dimer.^2^ Dimers can further organize into lattice-like oligomers, which may scaffold the viral envelope and contribute to membrane curvature during budding.

The multipass truncates 74mer and 107mer exhibit a strong tendency to oligomerize (**Figures 3E** and **S3**), with TMD2 being pivotal in orchestrating the association of the monomers. It interacts with TMD1 and TMD3 individually (**Figure 3C**), and increases the number of inter-monomer contacts per residue (**Figures 3G, 3H**, and **S4B**). TMD bundle oligomerization suggests co-translational oligomerization,^71^ a phenomenon that was supported when targeted mutation of four transmembrane residues disrupted both helical bundle assembly and downstream arrangement of the cytoplasmic domain (**Figures 4D** and **4E**). Detailed analysis of M folding pathway revealed two differentiated folding dynamics (**Figure 1C**): rapid TMD assembly, and delayed cytoplasmic β-sheet domain assembly. This suggests a potential regulatory mechanism in which the cytosolic portion remains partially unfolded until both monomers are available for stable dimer formation. A similar folding pathway for a multi-spanning membrane protein has been described in CFTR protein.^72,73^

Collectively, our data support a model in which M-protein dimers co-translationally assemble after TMD bundle insertion (**Figure 4E**), a process thought to be slower than the soluble domain folding.^74^

### Chaperone-assisted folding

After insertion, membrane proteins are assisted by chaperones to adopt their functional conformation and prevent aggregation.^75^ CANX and UGGT1 were the only other two genes that showed dependency for M stability in a CRISPRi screen.^32^ In our PL experiments, both proteins were identified in proximity to the 74mer and 107mer. The involvement of UGG glucosyltransferase 1 (UGGT1) might be especially interesting as it is a specific chaperone that corrects the folding of glycosylated unfolded domains,^76,77^ like the M-protein N-terminal region. Within the CANX and ER-chaperone clusters (**Figures 6E** and **6F**), we observed correlations with these two factors that revealed several additional chaperones and glycosylation-editing enzymes that may be promising targets for attenuating SARS-CoV-2 infection.

### Post-translational modifications

The strongest evidence for Sec61 translocon-independent insertion of TMD1 includes the post-translational N-terminal glycosylation of M by the OST-B complex.^47^ Our data reveal very low glycosylation efficiency for the 50mer variant (**Figure 2D**). We also observed strong labeling by 107mer of two OST-B-specific subunits, but not of the OST-A subunit STT3A or its adaptors (**Figure 5E**). Results consistent with OST-B’s role in post-translational glycosylation of proteins that bypass Sec61-mediated insertion.

### Blocking M-protein biogenesis as a therapeutic strategy

Recent studies identified two small-molecule inhibitors, CIM-834^7^ and JNJ-9676,^8^ which exhibit strong antiviral activity by blocking the M-protein transition from its extended to compact conformation. These findings demonstrate the therapeutic potential of targeting M protein biogenesis as an antiviral strategy.^78^ In this work, we identify new molecular events that may be targeted as new antiviral strategies.

Designing MPT and EMC inhibitors of specific subunits, such as EMC2 or TMCO1, could also be promising, as these components might not be essential for maintaining the integrity of the entire complex.^24,28^ Moreover, they are involved only in the insertion of a specific subset of multipass membrane proteins, potentially offering a safer therapeutic profile, compared to broad-spectrum Sec61 inhibitors.^79,80^ Additionally, targeting UGGT1 may destabilize the M N-terminal domain.^32^

We identified an specific cluster of hydrophobic residues critical for establishing inter-monomer hydrophobic contacts (**Figure 3B**). Mutations in this cluster disrupt oligomerization (**Figures 3B** and **3D**), impair co-translational folding (**Figure 3E**), reduce Spike protein recruitment (**Figure 3F**), and block VLP formation (**Figure 3G**).

Overall, the M protein is emerging as a promising antiviral target, particularly due to its high sequence conservation across sarbecoviruses, and its lower mutation rate compared to the Spike protein.^81^ By dissecting its insertion pathways and identifying critical host factors, we reveal multiple molecular targets that could be exploited to disrupt M-protein biogenesis and inhibit SARS-CoV-2 replication.

## Acknoledgements

We thank P. Selvi and E. Navarro for their excellent technical assistance. Certain cell culture and microscopy experiments were carried out in the Cell Culture and Flow Cytometry and the Microscopy facilities at the SCSIE (University of Valencia), respectively. LC-MS experiments were carried out at the Fracis Crick Institute Proteomics STP. This work was supported by grants ID2023-152568NB-I00 from the Spanish Ministry of Science, Innovation and Universities (MCIN/AEI/10.13039/501100011033) and CIPROM/2022/062 from the Generalitat Valenciana (to I.M.) by MCIN/AEI/https://doi.org/10.13039/501100011033 and European Union NextGenerationEU/PRTR. J.O-M. had a predoctoral contract granted by the University of Valencia’s “Atracció de Talent” program (UV-INV_PREDOC 1911519) and now has a research contract from CIPROM/2022/062 project (CPI-25-550). M.R-S is the recipient of a predoctoral contract from the Spanish Ministry of Science and Innovation (PRE2021-101042 by MCIN/AEI/10.13039/ 501100011033. J. A-R is a recipient of a predoctoral contract from the Spanish Ministry of Science, Innovation and Universities (FPU24/01943). J.G.C. is a Wellcome Trust Senior Research Fellow (224484/Z/21/Z). J.G.C. and M.S are supported by the Francis Crick Institute that receives its core funding from Cancer Research UK (CC2166,CC1063), the UK Medical Research Council (CC2166, CC1063), and the Wellcome Trust (CC2166, CC1063). For the purpose of Open Access, the authors have applied a CC BY public copyright licence to any Author Accepted Manuscript version arising from this submission. This work was supported by grants from the Knut and Alice Wallenberg Foundation (2017.0323), the Novo Nordisk Fund (NNF18OC0032828), and the Swedish Research Council (2020-03238) to G.v.H. J.C.G. was supported by the National Institutes of Health (R01-GM148586).

## Author contributions

J.O.-M. performed research except for the *pho-lac* dual reporter system, BlaTM and HiBiT assays, which were performed by J.M. A-C., M. R-S, and J. A-R., respectively. A. P. performed molecular dynamics simulations. G. J. P. provided experimental support and supervision for the proximity labeling experiments. S.S. provided experimental support for force profile analyses. A.M. and G.v.H. supervised the force profile analyses. J. M. S. supervised the LC-MS/MS experiments. J. C. G. supervised molecular dybnamics simulations. M.M. S. d.P. performed statistical analysis for proteomics data. J. O-M. performed data visualization. J. O-M., L. M.-G. and I. M. set the hypothesis and led the experimental design. I. M., L.M.-G. and J.O.-M. wrote the manuscript. G. v. H, J. G. C., J. C. G., I.M. and L.M.-G. secured funding.

## Materials and Methods

### Plasmid constructs

Sequence encoding for SARS-CoV-2 E, N and M protein (Gen Bank: QHD43419.1) were synthesized by ThermoFisher (GeneArtTM gene synthesis). A plasmid for SARS-CoV-2 Spike protein was a kind gift from Nevan Krogan. All four structural proteins were PCR amplified and subcloned into a KpnI-linearized pCAGGS vector. Different truncated DNA sequences from M gene were cloned with a c-myc epitope at their C-terminus, some of them adding a glycosylable NSTMMS or QSTMMS mock site. Alternatively, 74mer and 100mer truncates were added a longer linker (MGSMSGMGSMSGN/QSTMMS) to ensure that the putative glycosylation would be at least at a 11-residue distance from the membrane. M-SecM pING1 constructs were also synthesized by ThermoFisher and an N-terminal Lep (residues 1-99) tag was PCR amplified and fused to all constructs. TurboID constructs were generated by PCR amplifying M and Sec61β inserts and sub-cloning them into an HA-TurboID pCR3.1 vector upon HindIII/NotI restriction-ligation in case of C-terminal tagging, and EcoRI/NotI for N-terminal tagging. To generate BiFC chimeric plasmids including the Nt or Ct of the Venus Fluorescent Protein (VN, VC respectively; Addgene #27097, #22011, a gift from Chang-Deng H) plasmids were modified to clone the M protein truncates at the Nt of the VFP. For the VLP entry assay, sequences encoding M proteins tagged with the HiBiT peptide (VSGWRLFKKIS) in C terminus and LgBiT protein were PCR amplified and subcloned into a KpnI-linearized pCAGGS vector. All constructs were cloned using either In-Fusion HD cloning kit (Takara, Japan) or Gibson Assembly (NEB, USA) according to the manufacturer’s instructions. Site-directed mutagenesis was carried out using Pfu Plus! Polymerase (EurX, Poland) also according to manufacturer’s instructions. The substitutions included in the M TMD-Ala were designed to disrupt potential hydrophobic interactions, while preserving proper TMD insertion into the ER membrane. The sequences were verified by sequencing the plasmid DNA at Macrogen (South Korea), Azenta Life Biosciences (USA) or Eurofins Genomics (Germany).

### Force Profile analysis in *E. coli*

Force profile analysis (FPA) enables residue-level resolution of membrane integration events during translation.^33–36^ This technique quantifies the mechanical forces exerted on the nascent polypeptide chain while TMDs insert into the membrane and fold. This is achieved using the 17-residue *E. coli* SecM protein arrest peptide (AP) that binds in the ribosome exit tunnel, below the peptidyl transferase center, thereby stalling translation.^37^ If the nascent chain is subjected to a strong PF when the last AP codon is translated, the AP is removed from its binding site and translation proceeds at its normal rate.^38^ Therefore, the AP can be used as a “molecular force sensor” to probe co-translational force-generating processes, e.g., protein folding or membrane insertion. The M protein was truncated from its C-terminus at 5-residue resolution. Each construct was fused to the SecM AP, followed by a GGS linker and a C-terminal tail derived from *E. coli* LepB (residues 305– 324) can clone in a pING1 vector. We also fused LepB residues 1–99 (including its two TMDs) upstream of the M protein, to increase expression and ensure efficient targeting to the inner membrane SecYEG translocon. Competent E. coli MC106189 were transformed with the indicated constructs and grown overnight at 37°C in M9 minimal medium supplemented with 19 amino acids (1 mg/ml, no Met), 100 mg/ml thiamine, 0.4% (w/v) fructose, 100 mg/ml ampicillin, 2 mM MgSO4, and 0.1 mM CaCl2. Cells were diluted into fresh M9 medium to an OD600 of 0.1 and grown until an OD_600_ of 0.3–0.5. Expression from pING1 was induced with 0.2% (w/v) arabinose and continued for 5 min at 37°C. Proteins were then radiolabeled with [35S]-methionine for 10 min (2 min for constructs spanning the TMD region, 17-112) at 37°C before the reaction was stopped by adding ice-cold trichloroacetic acid (TCA) to a final concentration of 10%. Samples were put on ice for 30 min and precipitates were spun down for 10 min at 20,000 g at 4°C in a tabletop centrifuge (Eppendorf, Germany). After one wash with ice-cold acetone, centrifugation was repeated and pellets were subsequently solubilized in Tris-SDS buffer (10 mM Tris-Cl pH 7.5, 2% [w/v] SDS) for 5 min while shaking at 1000 rpm at 37°C. Samples were centrifuged for 5 min at 20,000 g to remove insoluble material. The supernatant was then added to a buffer containing 50 mM Tris-HCl pH 8.0, 150 mM NaCl, 0.1 mM EDTA-KOH, 2% (v/v) triton X-100, and supplemented Protein-G-agarose (Roche). After 15 min incubation on ice, non-specifically bound proteins were removed by centrifugation at 20,000 g. The supernatant was used for immuno-precipitation using Protein-G-agarose and Anti-HA.11 Epitope Tag Antibody (mouse) (BioLegend). The incubation was carried out at 4°C while rolling. After centrifugation for 1 min, immunoprecipitates were washed with 10 mM Tris-Cl pH 7.5, 150 mM NaCl, 2 mM EDTA, and 0.2% (v/v) triton X-100 and subsequently with 10 mM Tris-Cl pH 7.5. Samples were spun down again and pellets were solubilized in SDS sample buffer (67 mM Tris, 33% [w/v] SDS, 0.012% [w/v] bromophenol blue, 10 mM EDTA-KOH pH 8.0, 6.75% [v/v] glycerol, 100 mM DTT) for 10 min while shaking at 800 rpm. Solubilized proteins were incubated with 0.25 mg/ml RNase for 30 min at 37°C and subsequently separated by SDS-PAGE on Bis-Tris gels (Thermo Fisher Scientific). Gels were fixed in 30% (v/v) ethanol and 10% (v/v) acetic acid and dried by using a Bio-Rad gel dryer model 583 (Bio-Rad Laboratories, US). Radiolabeled proteins were detected by exposing dried gels to phosphorimaging plates, which were scanned in a Fujifilm FLA-3000 scanner (Fujifilm, Japan). Band intensity profiles were obtained using the FIJI (ImageJ) software and quantified with EasyQuant software. A and/or FL controls were included in the SDS-PAGE analysis for constructs where the identities of the A and FL bands were not immediately obvious on the gel. Data was generally collected from three independent biological replicates and mean ±SD were calculated.

### *Pho-lac* Dual reporter assay

The bacterial *pho-lac* dual-reporter assay for membrane proteins^42^ is based on two enzymatic reporters, alkaline phosphatase (PhoA) and β-galactosidase (LacZα fragment) fused to the protein of interest. When the chimeric membrane protein/*pho-lac* fusion is oriented with the C-terminus in the periplasm, PhoA becomes active, yielding high alkalinity and low LacZα activity, resulting in blue colonies. Conversely, if the fusion is oriented toward the cytoplasm, phosphatase activity is lost, while LacZ activity becomes prominent upon LacZα complementation with the soluble ω fragment, yielding red colonies. To quantify enzymatic activity, we used standard colorimetric substrates, allowing normalized activity ratio (NAR) calculation.^43^ DNA sequences encoding four protein truncates (50mer, 74mer, 100mer, 107mer) plus full-length M sequence were cloned in pKTop plasmid, each construct was transformed into competent E. coli DH5α cells and these were routinely grown at 37°C in agar plates with antibiotic 50 µg/mL kanamycin. To study the topology of the M protein in a qualitative assay, a freshly transformed colony was selected, the cells were grown up to OD600 of 0.5 in Luria-Bertani (LB) broth (0.5% yeast extract, 1% tryptone, 0.5% NaCl) and 7 µl was spotted onto plates contain Luria broth agar supplemented with the indicators 80 µg/mL disodium 5-bromo-4-chloro-3-indolyl phosphate (BCIP) and 100 µg/mL 6-chloro-3-indolyl-β-D-galactoside (Red-Gal), the inducer 1 mM IPTG, and the antibiotic 50 µg/mL kanamycin. Plates were developed overnight at 37 °C. In quantitative test, the cells were grown up to OD600 of 0.5 in LB with antibiotic 50 µg/mL kanamycin, at which cultures were induced with 1 mM IPTG for one hour with aeration at 37 °C to induce expression of truncates/pholac protein. To study β-galactosidase activity, the cells were harvested by centrifugation for 5 min at 7000 rpm and resuspended in M63 medium (100 mM KH2PO4, 15 mM (NH4)2SO4, 1.7 mM Fe2SO4, 1 mM MgSO4, pH 7.0). Then 50 µl of bacterial suspension, permeabilized with SDS/chloroform, was mixed with 100 µl of PM2 buffer and ONPG (0.15%) to initiate the enzymatic reaction. To study phosphatase activity, the cells were harvested by centrifugation for 5 min at 7000 rpm and resuspended in cold PM1 buffer (1 M Tris–HCl, pH 8.0, 0.1 mM ZnCl2, 1 mM iodoacetamide). Then 100 µl of bacterial suspension permeabilized with SDS/chloroform, was mixed with 50 µl of the pNPP solution (0.15% in 1 M Tris–HCl, pH 8.0) to initiate the enzymatic reaction. The β-galactosidase and phosphatase reactions were stopped by adding 1 M Na2CO3 and 2 N NaOH respectively and the enzymatic activities in relative units and the NAR was calculated as described.

### Cell lines and culture conditions

Wild-type human embryonic kidney (HEK) 293T and Vero E6 cells were acquired from American Type Cell Collection (ATCC, USA). A 293T cell line stably expressing human ACE2 and TMPRSS2 was acquired from BEI Resources (USA, catalog nº NR-55293). All lines were maintained in Dulbecco’s modified Eagle’s medium (DMEM, Gibco) supplemented with 10 % fetal bovine serum, 1 % penicillin-streptomycin (Sigma Aldrich, St. Louis, MO, USA), and 1 % MEM non-essential amino acids. Cells were maintained at 37°C and 5 % CO2 conditions.

### Transient DNA transfections

HEK293T or VeroE6 cells were seeded in 6-well, 12-well, 24-well plates, 15 cm disks or T75 flasks and grown in DMEM (Gibco) supplemented with 10 % fetal bovine serum (FBS, Gibco) the day before transfection. After 24 h, 100 µl of Op-tiMEM medium (Gibco) was mixed with plasmid DNA and either 3 µl PEI for 293T transfection or 1 µl Lipofectamine 3000 for VeroE6 transfections (per µg of DNA) and incubated for 15 minutes. For western-blotting, 4 × 10^6^ HEK 293T cells were transfected with 2 µg of the appropriate plasmids. For BiFC experiments, 500 ng cMyc-M-VN and 500 ng HA-M-VC were transfected into 2 × 10^6^ HEK 293T cells together with a plasmid expressing Renilla luciferase under the CMV promoter (pRL-CMV) (100 ng) in 12-well plates. For TurboID experiments, cells were transfected with either WT or truncate M-HA-TurboID, Sec61β-HATurboID, HA-TurboID-Sec61β and mEmerald-HA-TurboID constructs using PEI, with 1800 µL optiMEM, 36 µg DNA, and 72 µL PEI used per T75 flask. For HiBiT-tagged VLP entry assays, 2 x 10^6^ HEK 293T cells in 6-well plates were transfected with 2 µg M-HiBiT and 2 µg S (producer cells). On the other hand, 1 x 10^6^ HEK 293T cells in 12-well plates were transfected with 250 ng LgBiT and 500 ng hAce2 (receptor cells). In the BiMuC assay, 2 × 10^6^ HEK 293T cells in 6-well plates were transfected with 1 µg of S, 1 µg of VN-Jun and 1 µg of M-cMyc. Alternatively, 293T-hAce2-TMPRSS2 cells were transfected with 1 µg of VC-Fos, plus 100 ng pRL-CMV per µg of DNA. For VLP production, pCAGGS plasmids coding for SARS-CoV-2 structural M-cMyc plasmids, S, E and N proteins (9 µg of each plasmid, up to a total of 36 µg) and transfected into 150 mm disks. Trans-fection mixtures were added dropwise onto cells and media was changed after 6h.

### Protein expression and western blotting

Two days after transfection, media was aspirated, cells rinsed with PBS and lysed in 300 µl of radioimmunoprecipitation assay (RIPA) buffer [150 mM NaCl, 50 mM tris-HCl (pH 8), 1% NP-40, 0.5 % sodium deoxycholate, 0.4% SDS, and 1 mM EDTA] supplemented with cOmplete EDTAfree protease inhibitors (Roche). Lysates were sonicated with three pulses of 1 s in a VCX-500 Vibra Cell sonicator (Sonics) following addition of 5X sample buffer (final concentration 62.5 mM Tris-HCl [pH 6.8], 2 % sodium dodecyl sulfate [SDS], 0.01 % bromophenol blue, 10 % glycerol and 5 % β-mercaptoethanol). Protein samples were boiled for 4 min at 95 ºC, subjected to 10 % SDS-polyacrylamide gel electrophoresis (PAGE), and transferred to nitrocellulose membranes (Cytiva). Membranes were blocked for 30 min at room temperature in Tris-buffered saline supplemented with 0.05 % Tween 20 (TBS-T) containing 5% non-fat dry milk and later incubated with primary antibodies diluted in the same buffer at 4 °C overnight. Antibodies used in this study were rabbit anti-SARS-CoV-2 M (GeneTex GTX636246), rabbit anti c-Myc (Sigma PLA0001), mouse anti-HA.11 (Biolegend) and mouse anti-GAPDH (Santa Cruz sc-47724). Then, membranes were washed with TBS-T and incubated with goat anti-rabbit or sheep anti-mouse IgG horseradish peroxidase conjugate (Sigma DC02L or GE Healthcare NXA931, respectively) for 1 h at room temperature and washed again. All antibodies were diluted in TBS-T with 5 % non-fat dry milk. Detection of immunoreactive proteins was carried out using the enhanced chemiluminescence reaction (Amersham ECL Prime, Cytiva) and detected by the Image Quant LAS 4000 Mini (GE Healthcare).

### Confocal microscopy and immunofluorescence

Vero E6 cells (2 × 10^3^ cells/well) were seeded on 10 mm coverslides in 24-well plates. The next day, cells were transfected with the appropriate TurboID or M-cMyc plasmids. To limit overexpression, cells were fixed at 16 hours post-transfection with 4% paraformaldehyde for 20 min, then washed 3 times with PBS. For immunolabelling against HA, cMyc tags or endogenous STT3B, cells were permeabilized with 0.1 % Triton-X100 in PBS, washed 3 times in PBS, and blocked in 1% BSA for 1 hour. After primary and secondary antibody incubations, coverslips were mounted on X50 SuperFrost microscope slides using Mowiol. Imaging was performed using an Andor Dragonfly 200 spinning disc confocal paired with a Zyla 5.5 sCMOS camera and using a Nikon Eclipse Ti2 with Plan Apo 60×/1.4 numerical aperture (NA). Laser intensity was individually adjusted in all samples.

### BLaTM assay

To assess TMD–TMD interactions we used a BLaTM in *E. coli*. In this assay the TMDs of interest are fused to the N-or C-terminus of split β-lactamase fragments (βN and βC). Productive TMD–TMD interactions facilitate β-lactamase enzyme reconstitution, enabling bacterial growth on selective media. Given the M protein’s multipass nature, we applied two complementary strategies to test parallel and antiparallel interactions.^82^ In the parallel configuration glycophorin A (GpA), a membrane-spanning helix that forms noncovalent dimers, was used as a positive control and normalization reference.^83–85^ Negative controls included a 20-residue poly-leucine stretch (Leu20) and the non-oligomerizing C-terminal TMD of monomeric L-selectin (CLS).^86,50^ In the antiparallel configuration, we used the fourth TMD of the *EmrE* bacterial protein (an antiparallel dimer) as a positive control, and an interaction-deficient double mutant of the same TMD (G90V/G97V, *EmrE* mut) as a negative control, alongside Leu20.^82^ Competent E. coli BL21-DE3 cells were co-transformed with N-BLa and C-BLa plasmids, version 1.2 and 2.0,^82^ containing a given TMD pair and grown overnight at 37 °C on LB-agar plates containing 34 µg/mL of chloramphenicol (Cm) and 35 µg/mL of kanamycin (Kan) for plasmid inheritance. After o/n incubation at 37 °C, colonies were either picked for immediate use or the plates were sealed with Parafilm (Pechiney Plastic Packaging) and stored at 4 °C for up to one week. Overnight cultures were conducted by inoculating 5 mL of LB-medium (Cm, Kan) with 10 colonies from one agar plate, followed by o/n incubation in an orbital incubator at 37 °C, 200 rpm. An expression culture was started with a 1:10 dilution of the overnight culture in 4 mL expression medium: LB medium (Cm, Kan) containing 1.33 mM arabinose. After 4 h at 37 °C, the expression cultures were diluted to an OD600 = 0.1 in the expression medium. To expose the bacteria to different ampicillin concentrations, an LD50 culture was prepared by pipetting 100 µL of the diluted expression culture into each cavity of a 96-deep well plate (96 square well, 2 mL, VWR) containing 400 µL of expression media (final OD600 = 0.02). Freshly prepared ampicillin stock (100 mg/mL in ethanol) was added, resulting in ampicillin concentrations ranging up to 350 µg/mL, depending on the affinity of the TMD under investigation. As a rule, the maximum ampicillin concentration to be used for a particular case should be about twice the mean LD50. The plates were incubated in a moisturized container for 16 h at 37 °C and 250 rpm on a shaker (shaking amplitude 10 mm, KS 260 Basic, IKA) containing tips in every well to ensure proper agitation. Cell density was measured via absorbance at 544 nm in a microplate reader (Victor X3, Perkin Elmer). To minimize clonal variation, at least two transformations were done and at least two separate LD50 cultures were inoculated from each batch of transformed bacteria using ten colonies for each culture. Thus, at least 40 colonies entered each determination of LD50.

### Bimolecular fluorescent complementation (BiFC) assay

Two non-fluorescent fragments of the Venus fluorescent protein (VN and VC)^87^ were fused to the 50mer, 74mer, and 107mer variants, and to the full-length M protein, to assess whether interaction between the tested polypeptides restored fluorescence. Then, 48 h after transfection, PBS washed and collected for fluorescence and luciferase measurements (Victor X3 plate reader). For the Renilla luciferase readings used as signal normalization, we used the Renilla Luciferase Glow Assay Kit (Pierce, Thermofisher) according to the manufacturer’s protocol. In each experiment, the fluorescence/luminescence ratio obtained with the M-M homo-oligomer was used as a 100% oligomerization value and the rest of the values adjusted accordingly. All experiments were done at least in triplicates.

### Molecular dynamics simulations

The starting structure for the M protein dimer was generated with AlphaFold2^88^ before the crystal structures were released. The monomer and/or truncated constructs were all generated from this structure by removing relevant residues. All systems were embedded in symmetric membranes using CHARMM-GUI.^89^ The ER membrane was simulated using 54 POPC, 20 POPE, and 11 POPI lipids, as well as 8 cholesterol molecules in each leaflet. The membrane-embedded proteins were solvated, and NaCl was added at 0.15 M concentration. All simulations were performed with NAMD3^90^ using the CHARMM36m^91^ forcefield and TIP3P water model.^92^ In all simulations we used rigid bonds for covalent hydrogen bonds, which allowed us to integrate the equations of motion with a 2-fs time step. Van der Waals interactions were cut off at 12 Å, and a smoothing function was applied from 10 to 12 Å to ensure a smooth decay to zero. Long-range electrostatic interactions were calculated using the particle-mesh Ewald method.^93^ The temperature and the pressure were kept constant at the biologically relevant values of 310 K and 1 bar, respectively, with a Langevin piston barostat and Langevin thermostat. For the Langevin piston we used a period of 300 fs and a decay of 150 fs. Equilibration of systems was done in several 500-ps-long steps: first the position of all hydrogen atoms was minimized, followed by equilibration of lipid tails, restraining the rest of the system. Next, water, ions and the membrane were equilibrated while restraining the protein; in the final equilibration step everything except the protein backbone was released. After equilibration, 400-ns-long simulations were performed for each system in triplicate.

### HiBiT-tagged VLP entry assay

At 3 hours after transfection, receptor cells’ media was discarded and 1 mL DMEM was added to each well. Cells were collected, counted and 100 µL containing 1 x 10^4^ cells were seeded into a 96-white-well plate. Two days post transfection, producer cells’ media was collected and centrifuged for 10 min at 300 g. Supernatant was saved. Alternatively, receptor cells’ media was discarded, cells rinsed with PBS and 200 µL of the supernatant from the producer cells were added to each well. Following a 3-hour incubation, media was discarded, 50 µL PBS containing 1 µM DrkBiT peptide^94^ (VSGWALFKKIS, custom-synthetized by GenScript, USA) was added and, after the addition of 10 µL of the Nano-Glo® Live Cell Assay System (Promega), luminescence was measured in a VictorX multi-plate reader (Perking Elmer, USA). RLU values were then normalized against the wild-type condition.

### Bimolecular Multicellular Complementation (BiMuC) assay

One day post-transfection, media was discarded and cells rinsed with PBS. Then, 1 mL DMEM was added to each well and one well expressing S and/or M and VN-Jun was pooled with other well expressing VC-Fos in an hAce2-TMPRSS2 background.^95^ Cells were counted and 500 µl containing 3 x 10^5^ cells were seeded into 24-well plates in triplicates. The next day, media was discarded, cells rinsed with PBS, collected in 100 µl PBS, and seeded into 96-well black plates for fluorescence read. Fluorescence was measured in a VictorX multi-plate reader (Perking Elmer, USA). The luminescence was measured using the Renilla Luciferase Assay kit from Sigma following the manufacturer’s protocol on 96-well white cell-culture plates. Five independent experiments were done. The ratio between VFP and luciferase was registered as the relative fluorescence units (RFU) for each well. Each of the three wells were accounted as technical replicates, and the mean of the three values was taken as a biological replicate. RFU values were then normalized against the wild-type condition.

### VLP production

At 48 hours after transfection, media was collected from 150 mm dishes and centrifuged at 1000 g. Supernatant was cleared with 0.45 µm filters and transferred to a 13.2 mL ultracentrifuge tube before the addition of 1 mL of a 20% sucrose cushion with a 12 cm syringe. Media samples were then centrifuged for 2 h at 28000 rpm using a SW Ti 41 rotor in an Optima XE ultracentrifuge (Beckman Coulter, USA). Supernatant was discarded and pellets were resuspended in 50 µl of PBS and processed for western blot analysis. In parallel, cells from the culture disk were collected by centrifugation and lysed for western blot analysis as described previously.

### TurboID Proximity Biotinylation and Pull Down

To identify cellular factors involved in M-protein insertion and folding in the ER, we investigated the local environment during its biogenesis. We employed a proximity labeling strategy, fusing an HA-TurboID tag to the full-length protein and the 50mer, 74mer, and 107mer variants. TurboID is an engineered biotin ligase with high temporal resolution, which enables labeling of proteins within a ~10-nm radius in live cells.^96,97^ Several studies have used proximity labeling to identify SARS-CoV-2 host partners using M protein as bait.^98–100^ We aimed to define and compare the proximal proteomes of full-length and truncated M variants, to obtain snapshots of the biogenesis process. We tagged the C-terminal end of all constructs, meaning that in the 50mer, 107mer, and full-length M constructs, TurboID faced the cytosol (C^cyt^ orientation). The 74mer construct translocates the TurboID tag into the ER lumen upon insertion of the second TMD, yielding a luminally oriented receptor (C^exo^). The cytosolic mEmerald fluorescent protein served as a non-membrane control. We also used the tail-anchored Sec61β subunit of the ER translocon, fused to TurboID at its N-terminus (cytosolic orientation, Sec^cyt^) or C-terminus (luminal orientation, Sec^exo^). Adding a large C-terminal tag to a tail-anchored protein may alter its membrane insertion pathway, shifting from Oxa1-dependent to Sec61-mediated insertion; however, a similar construct has reportedly maintained its functional topology *in vivo*.^101^ These controls allowed us to differentiate true partners from background signals, confirm labeling of known ER-associated proteins, and validate luminal versus cytosolic orientation of the TurboID tag. Comparing the 74mer construct with Sec^exo^ provided insight regarding luminal partners specific to M’s membrane-embedded regions. HEK 293-T cells were seeded to T75 flasks, with 3 flasks per condition. Exactly 18 h after transfection, cells were biotinylated by incubation for 20 minutes in biotin-free growth media supplement with 50 µM biotin (Sigma-Aldrich). At 20 minutes exactly, flasks were placed on ice and washed 1x with ice-cold PBS to halt the biotinylation reaction. Cells were scrapped in 10 ml of ice-cold PBS and pelleted, with pellets kept on ice until all samples had been prepared. Cell pellets were then lysed by 30 min incubation at 4ºC in 1 ml RIPA buffer (150 mM NaCl, 50 mM Tris-HCl pH8, 1% NP40, 0.5% Sodium deoxycholate, 0.4% SDS, 1 mM EDTA) supplemented with complete EDTA-free protease inhibitors (Roche) and 167 U/ml of Benzonase Nuclease (Sigma-Aldrich). During this time preacetylated NeutrAvidin agarose beads (Pierce) were washed 4 times with 10x their volume in lysis buffer, with 40 µL beads used per sample. NeutrAvidin bead acetylation was necessary to stop the Neutravidin being cleaved from the agarose beads during on-bead digestion and performed prior to the day of pulldown by two 30 min incubations of beads with 10 mM Sulfo-NHS acetate (ThermoFisher) on a rotating wheel followed by quenching in 90 mM Tris-HCl pH7.5. Lysed cell pellets were centrifuged at 14,000 rpm at 4ºC for 15 mins to sediment undigested nuclear debris, and lysate supernatants were mixed with equal amounts of washed acetylated NeutrAvidin beads and rotated at room temperature for 2 hrs. The beads were then washed 2 times in 500 uL RIPA buffer and 5 times in 25 mM HEPES pH 8.5, with the beads rotated for 5 mins at room temperature for each wash. After the final wash, beads were resuspended in 100 µL 25 mM HEPES pH 8.5 and 100 ng of Lysyl endopeptidase LysC (WAKO) was added to each sample, with this mixture incubated for 16 h at 37ºC in a hooded ThermoMixer at 1,200 rpm. Each bead supernatant was then transferred to a new Eppendorf and mixed with 100 ng Trypsin (Pierce) and incubated at 37ºC for 6 h. The solutions were then acidified to a final concentration of 0.5% trifluoroacetic acid (TFA).

### LC-MS/MS analysis

The resulting peptides were analysed by nano-scale capillary LC-MS/MS using an Ultimate U3000 HPLC (ThermoScientific Dionex, San Jose, USA) to deliver a flow of approximately 250 nL/min. A C18 Acclaim PepMap100 5 µm, 75 µm x 20 mm nanoViper (ThermoScientific Dionex, San Jose, USA), trapped the peptides prior to separation on a C18 Acclaim PepMap RSLC 3 µm, 75 µm x 500 mm nanoViper (ThermoScientific Dionex, San Jose, USA). Peptides were eluted with a 90 min gradient of acetonitrile (2%v/v to 80%v/v). The analytical column outlet was directly interfaced via a nano-flow electrospray ionisation source, with a hybrid dual pressure linear ion trap mass spectrometer (Orbitrap Lumos, Thermo-Scientific, San Jose, USA). Data dependent analysis was carried out, using a resolution of 120,000 for the full MS spectrum, followed by ten MS/MS spectra in the linear ion trap. MS spectra were collected over a m/z range of 275–1500. MS/MS scans were collected with the standard AGC target, dynamic maximum injection time mode, isolation window at 1.2 m/z and 32% normalised HCD collision energy. All raw files were processed with MaxQuant 1.6.12.0^102^ using standard settings and searched against a UniProt Human Reviewed KB database with the Andromeda search engine^103^ integrated into the MaxQuant software suite. Enzyme search specificity was Trypsin/P. Up to two missed cleavages for each peptide were allowed. Carbamidomethylation of cysteines was set as fixed modification with oxidized methionine and protein N-acetylation considered as variable modifications. The search was performed with an initial mass tolerance of 6 ppm for the precursor ion and 0.5 Da for MS/MS spectra. The false discovery rate was fixed at 1% at the peptide and protein level. Statistical analysis was carried out using the Perseus module (v1.4.0.2) of MaxQuant. Prior to statistical analysis, peptides mapped to known contaminants, reverse hits and protein groups only identified by site were removed. Only protein groups identified with at least two peptides, one of which was unique and two quantitation events were considered for data analysis.

### Proteomics data analysis

Before processing the quantitative data provided by MaxQuant, contaminating (Crapome V2.39) and decoy proteins were removed, as well as proteins that were not detected at least twice in a technical replicate. Preliminary inspection of the data revealed that the number of proteins analyzed was abnormally low in some LC-MS/MS runs. These runs were considered outliers (fewer than 1,400 identified proteins) and, therefore, were not included in the analysis. Finally, only proteins detected in 80% of at least one sample type were considered in the analysis. Quantitative values were log2 transformed, median normalized. P-values were calculated by two-sample t-tests using Benjamini-Hochberg FDR correction for multiple testing. It is to be expected that spatially close proteins have similar labeling patterns. To determine the similarity of the labeling patterns, a correlation analysis was performed by calculating Pearson’s coefficients among proteins. Only those coefficients calculated with at least 50 data points were taken into consideration. Positive correlations with coefficients greater than 0.8 were transferred to Cytoscape^104^ to generate protein networks with similar labeling patterns. A few protein clusters were clearly apparent and were further clustered using the MCL algorithm of clusterMaker2.^105^ Clusters were functionally analyzed using Cytoscape’s stringApp.^106^

### Sequence alignments

Alignments of coronavirus Membrane sequences were performed using T-Coffee.^107^ Aligned sequences were then exported and viewed in Jalview^108^, and residues were color-coded using ClustalX color map.

### Statistical analysis

Unpaired t-tests assuming Gaussian distribution and equal standard deviation (SD) for all conditions were applied. The p-value was two-tailed. One sample t-test was performed to compare different wild-type-normalized conditions, comparing means with a ‘hypothetical value’ of 100. Significance assessment between test samples and controls was performed using GraphPad Prism for P values < 0.05.

### Data availability

All data needed to evaluate the conclusions in the paper are present in the paper and/or the Supplementary Materials. Mass spectrometry proteomics data have been deposited to the ProteomeXchange consortium with the dataset identifier PXD064009.

## References

1. Boson, B., Legros, V., Zhou, B., Siret, E., Mathieu, C., Cosset, F.-L., Lavillette, D., and Denolly, S. (2021). The SARS-CoV-2 envelope and membrane proteins modulate maturation and retention of the spike protein, allowing assembly of virus-like particles. J. Biol. Chem. 296, 100111. 10.1074/jbc.RA120.016175.

2. Zhang, Z., Nomura, N., Muramoto, Y., Ekimoto, T., Uemura, T., Liu, K., Yui, M., Kono, N., Aoki, J., Ikeguchi, M., et al. (2022). Structure of SARS-CoV-2 membrane protein essential for virus assembly. Nat. Commun. 13, 4399. 10.1038/s41467-022-32019-3.

3. Zhang, Y., Anbir, S., McTiernan, J., Li, S., Worcester, M., Mishra, P., Colvin, M.E., Gopinathan, A., Mohideen, U., Zandi, R., et al. (2024). Synthesis, insertion, and characterization of SARS-CoV-2 membrane protein within lipid bilayers. Sci. Adv. 10, eadm7030. 10.1126/sciadv.adm7030.

4. Dolan, K.A., Dutta, M., Kern, D.M., Kotecha, A., Voth, G.A., and Brohawn, S.G. (2022). Structure of SARS-CoV-2 M protein in lipid nanodiscs. eLife 11, e81702. 10.7554/eLife.81702.

5. Neuman, B.W., Kiss, G., Kunding, A.H., Bhella, D., Baksh, M.F., Connelly, S., Droese, B., Klaus, J.P., Makino, S., Sawicki, S.G., et al. (2011). A structural analysis of M protein in coronavirus assembly and morphology. J. Struct. Biol. 174, 11–22. 10.1016/j.jsb.2010.11.021.

6. Dutta, M., Dolan, K.A., Amiar, S., Bass, E.J., Sultana, R., Voth, G.A., Stahelin, R.V., and Brohawn, S.G. (2024). Direct lipid interactions control SARS-CoV-2 M protein conformational dynamics and virus assembly. Preprint at bioRxiv, 10.1101/2024.11.04.620124 https://doi.org/10.1101/2024.11.04.620124.

7. Laporte, M., Jochmans, D., Bardiot, D., Desmarets, L., Debski-Antoniak, O.J., Mizzon, G., Abdelnabi, R., Leyssen, P., Chiu, W., Zhang, Z., et al. (2025). A coronavirus assembly inhibitor that targets the viral membrane protein. Nature, 1–10. 10.1038/s41586-02508773-x.

8. Van Damme, E., Abeywickrema, P., Yin, Y., Xie, J., Jacobs, S., Mann, M.K., Doijen, J., Miller, R., Piassek, M., Marsili, S., et al. (2025). A small-molecule SARS-CoV-2 inhibitor targeting the membrane protein. Nature, 1–8. 10.1038/s41586-025-08651-6.

9. Saurí, A., Tamborero, S., Martínez-Gil, L., Johnson, A.E., and Mingarro, I. (2009). Viral Membrane Protein Topology Is Dictated by Multiple Determinants in Its Sequence. J. Mol. Biol. 387, 113–128. 10.1016/j.jmb.2009.01.063.

10. Akopian, D., Shen, K., Zhang, X., and Shan, S. (2013). Signal Recognition Particle: An Essential Protein-Targeting Machine. Annu. Rev. Biochem. 82, 693–721. 10.1146/annurev-biochem072711-164732.

11. Duart, G., García-Murria, M.J., and Mingarro, I. (2021). The SARS-CoV-2 envelope (E) protein has evolved towards membrane topology robustness. Biochim. Biophys. Acta BBA - Biomembr. 1863, 183608. 10.1016/j.bbamem.2021.183608.

12. Jomaa, A., Fu, Y.-H.H., Boehringer, D., Leibundgut, M., Shan, S., and Ban, N. (2017). Structure of the quaternary complex between SRP, SR, and translocon bound to the translating ribosome. Nat. Commun. 8, 15470. 10.1038/ncomms15470.

13. Kobayashi, K., Jomaa, A., Lee, J.H., Chandrasekar, S., Boehringer, D., Shan, S., and Ban, N. (2018). Structure of a prehandover mammalian ribosomal SRP· SRP receptor targeting complex. Science 360, 323–327. 10.1126/science.aar7924.

14. Berg, B. van den, Clemons, W.M., Collinson, I., Modis, Y., Hartmann, E., Harrison, S.C., and Rapoport, T.A. (2004). X-ray structure of a protein-conducting channel. Nature 427, 36–44. 10.1038/nature02218.

15. Jaskolowski, M., Jomaa, A., Gamerdinger, M., Shrestha, S., Leibundgut, M., Deuerling, E., and Ban, N. (2023). Molecular basis of the TRAP complex function in ER protein biogenesis. Nat. Struct. Mol. Biol. 30, 770–777. 10.1038/s41594-023-00990-0.

16. Gemmer, M., Chaillet, M.L., van Loenhout, J., Cuevas Arenas, R., Vismpas, D., Gröllers-Mulderij, M., Koh, F.A., Albanese, P., Scheltema, R.A., Howes, S.C., et al. (2023). Visualization of translation and protein biogenesis at the ER membrane. Nature 614, 160–167. 10.1038/s41586-022-05638-5.

17. Gemmer, M., Chaillet, M.L., and Förster, F. (2024). Exploring the molecular composition of the multipass translocon in its native membrane environment. Life Sci. Alliance 7. 10.26508/lsa.202302496.

18. Lewis, A.J.O., Zhong, F., Keenan, R.J., and Hegde, R.S. (2024). Structural analysis of the dynamic ribosome-translocon complex. eLife 13. 10.7554/eLife.95814.2.

19. Sundaram, A., Li, Q., Wan, Y., Tang, J., Wu, H., Smalinskaitė, L., Hegde, R.S., Ji, Z., and Keenan, R.J. (2025). Global analysis of translocon remodeling during protein synthesis at the ER. Nat. Struct. Mol. Biol., 1–9. 10.1038/s41594-025-01691-6.

20. Guna, A., Volkmar, N., Christianson, J.C., and Hegde, R.S. (2018). The ER membrane protein complex is a transmembrane domain insertase. Science 359, 470–473. 10.1126/science.aao3099.

21. Chitwood, P.J., Juszkiewicz, S., Guna, A., Shao, S., and Hegde, R.S. (2018). EMC Is Required to Initiate Accurate Membrane Protein Topogenesis. Cell 175, 1507-1519.e16. 10.1016/j.cell.2018.10.009.

22. O’Donnell, J.P., Phillips, B.P., Yagita, Y., Juszkiewicz, S., Wagner, A., Malinverni, D., Keenan, R.J., Miller, E.A., and Hegde, R.S. (2020). The architecture of EMC reveals a path for membrane protein insertion. eLife 9, e57887. 10.7554/eLife.57887.

23. Chen, Z., Mondal, A., Abderemane-Ali, F., Jang, S., Niranjan, S., Montaño, J.L., Zaro, B.W., and Minor, D.L. (2023). EMC chaperone–CaV structure reveals an ion channel assembly intermediate. Nature 619, 410–419. 10.1038/s41586-023-06175-5.

24. Wu, H., and Hegde, R.S. (2023). Mechanism of signal-anchor triage during early steps of membrane protein insertion. Mol. Cell 83, 961-973.e7. 10.1016/j.molcel.2023.01.018.

25. Smalinskaitė, L., and Hegde, R.S. (2023). The Biogenesis of Multipass Membrane Proteins. Cold Spring Harb. Perspect. Biol. 15, a041251. 10.1101/cshperspect.a041251.

26. McGilvray, P.T., Anghel, S.A., Sundaram, A., Zhong, F., Trnka, M.J., Fuller, J.R., Hu, H., Burlingame, A.L., and Keenan, R.J. (2020). An ER translocon for multi-pass membrane protein biogenesis. eLife 9, e56889. 10.7554/eLife.56889.

27. Sundaram, A., Yamsek, M., Zhong, F., Hooda, Y., Hegde, R.S., and Keenan, R.J. (2022). Substrate-driven assembly of a translocon for multipass membrane proteins. Nature 611, 167–172. 10.1038/s41586-022-05330-8.

28. Smalinskaitė, L., Kim, M.K., Lewis, A.J.O., Keenan, R.J., and Hegde, R.S. (2022). Mechanism of an intramembrane chaperone for multipass membrane proteins. Nature 611, 161–166. 10.1038/s41586-022-05336-2.

29. Sundaram, A., Li, Q., Wan, Y., Tang, J., Wu, H., Smalinskaitė, L., Hegde, R.S., Ji, Z., and Keenan, R.J. (2025). Global analysis of trans-locon remodeling during protein synthesis at the ER. Nat. Struct. Mol. Biol., 1–9. 10.1038/s41594-025-01691-6.

30. Chitwood, P.J., and Hegde, R.S. (2020). An intramembrane chaperone complex facilitates membrane protein biogenesis. Nature 584, 630– 634. 10.1038/s41586-020-2624-y.

31. Meacock, S.L., Lecomte, F.J.L., Crawshaw, S.G., and High, S. (2002). Different Transmembrane Domains Associate with Distinct Endoplasmic Reticulum Components during Membrane Integration of a Polytopic Protein. Mol. Biol. Cell 13, 4114–4129. 10.1091/mbc.e02-04-0198.

32. Page, K.R., Nguyen, V.N., Pleiner, T., Tomaleri, G.P., Wang, M.L., Guna, A., Hazu, M., Wang, T.-Y., Chou, T.-F., and Voorhees, R.M. (2024). Role of a holo-insertase complex in the biogenesis of biophysically diverse ER membrane proteins. Mol. Cell 84, 3302-3319.e11. 10.1016/j.molcel.2024.08.005.

33. Ismail, N., Hedman, R., Schiller, N., and von Heijne, G. (2012). A biphasic pulling force acts on transmembrane helices during translocon-mediated membrane integration. Nat. Struct. Mol. Biol. 19, 1018–1022. 10.1038/nsmb.2376.

34. Nicolaus, F., Metola, A., Mermans, D., Liljenström, A., Krc, A., Abdullahi, S.M., Zimmer, M., Miller III, T.F., and von Heijne, G. (2021). Residue-by-residue analysis of cotranslational membrane protein integration in vivo. eLife 10, e64302. 10.7554/eLife.64302.

35. Nilsson, O.B., Hedman, R., Marino, J., Wickles, S., Bischoff, L., Johansson, M., Müller-Lucks, A., Trovato, F., Puglisi, J.D., O’Brien, E.P., et al. (2015). Cotranslational Protein Folding inside the Ribosome Exit Tunnel. Cell Rep. 12, 1533–1540. 10.1016/j.celrep.2015.07.065.

36. Westerfield, J.M., Kozojedová, P., Juli, C., Metola, A., and von Heijne, G. (2025). Cotranslational membrane insertion of the voltage-sensitive K+ channel KvAP. Proc. Natl. Acad. Sci. 122, e2412492122. 10.1073/pnas.2412492122.

37. Gersteuer, F., Morici, M., Gabrielli, S., Fujiwara, K., Safdari, H.A., Paternoga, H., Bock, L.V., Chiba, S., and Wilson, D.N. (2024). The SecM arrest peptide traps a pre-peptide bond formation state of the ribosome. Nat. Commun. 15, 2431. 10.1038/s41467-024-46762-2.

38. Goldman, D.H., Kaiser, C.M., Milin, A., Righini, M., Tinoco, I., and Bustamante, C. (2015). Mechanical force releases nascent chain–mediated ribosome arrest in vitro and in vivo. Science 348, 457–460. 10.1126/science.1261909.

39. Hessa, T., Kim, H., Bihlmaier, K., Lundin, C., Boekel, J., Andersson, H., Nilsson, I., White, S.H., and von Heijne, G. (2005). Recognition of transmembrane helices by the endoplasmic reticulum translocon. Nature 433, 377–381. 10.1038/nature03216.

40. Hessa, T., Meindl-Beinker, N.M., Bernsel, A., Kim, H., Sato, Y., Lerch-Bader, M., Nilsson, I., White, S.H., and von Heijne, G. (2007). Molecular code for transmembrane-helix recognition by the Sec61 translocon. Nature 450, 1026–1030. 10.1038/nature06387.

41. von Heijne, G. (1986). The distribution of positively charged residues in bacterial inner membrane proteins correlates with the trans-membrane topology. EMBO J. 5, 3021–3027. 10.1002/j.1460-2075.1986.tb04601.x.

42. Alexeyev, M.F., and Winkler, H.H. (1999). Membrane topology of the Rickettsia prowazekii ATP/ADP translocase revealed by novel dual pho-lac reporters11Edited by G. von Heijne. J. Mol. Biol. 285, 1503– 1513. 10.1006/jmbi.1998.2412.

43. Karimova, G., and Ladant, D. (2017). Defining Membrane Protein To-pology Using pho-lac Reporter Fusions. In Bacterial Protein Secretion Systems: Methods and Protocols Methods in Molecular Biology., L. Journet and E. Cascales, eds. (Springer), pp. 129–142. 10.1007/978-1-4939-7033-9_10.

44. Acosta-Cáceres, J.M., Lolicato, F., Gadea-Salom, L., Ortiz-Mateu, J., Belda, D., Martínez-Gil, L., García-Murria, M.J., and Mingarro, I. (2025). Structural insights into SARS-CoV-2 nonstructural protein 4 (nsp4) biogenesis. Protein Sci. 34, e70355. 10.1002/pro.70355.

45. Tamborero, S., Vilar, M., Martínez-Gil, L., Johnson, A.E., and Mingarro, I. (2011). Membrane Insertion and Topology of the Translocating Chain-Associating Membrane Protein (TRAM). J. Mol. Biol. 406, 571– 582. 10.1016/j.jmb.2011.01.009.

46. Bañó-Polo, M., Baldin, F., Tamborero, S., Marti-Renom, M.A., and Mingarro, I. (2011). N-glycosylation efficiency is determined by the distance to the C-terminus and the amino acid preceding an Asn-Ser-Thr sequon. Protein Sci. 20, 179–186. 10.1002/pro.551.

47. Juckel, D., Desmarets, L., Danneels, A., Rouillé, Y., Dubuisson, J., and Belouzard, S. (2023). MERS-CoV and SARS-CoV-2 membrane proteins are modified with polylactosamine chains. J. Gen. Virol. 104, 001900. 10.1099/jgv.0.001900.

48. Schanzenbach, C., Schmidt, F.C., Breckner, P., Teese, M.G., and Langosch, D. (2017). Identifying ionic interactions within a membrane using BLaTM, a genetic tool to measure homo- and heterotypic transmembrane helix-helix interactions. Sci. Rep. 7, 43476. 10.1038/srep43476.

49. Duart, G., Grau, B., Mingarro, I., and Martinez-Gil, L. (2021). Methodological approaches for the analysis of transmembrane domain interactions: A systematic review. Biochim. Biophys. Acta BBA - Biomembr. 1863, 183712. 10.1016/j.bbamem.2021.183712.

50. García-Murria, M.J., Duart, G., Grau, B., Diaz-Beneitez, E., Rodríguez, D., Mingarro, I., and Martínez-Gil, L. (2020). Viral Bcl2s’ transmembrane domain interact with host Bcl2 proteins to control cellular apoptosis. Nat. Commun. 11, 6056. 10.1038/s41467-020-19881-9.

51. Lucendo, E., Sancho, M., Lolicato, F., Javanainen, M., Kulig, W., Leiva, D., Duart, G., Andreu-Fernández, V., Mingarro, I., and Orzáez, M. (2020). Mcl-1 and Bok transmembrane domains: Unexpected players in the modulation of apoptosis. Proc. Natl. Acad. Sci. 117, 27980– 27988. 10.1073/pnas.2008885117.

52. Jumper, J., Evans, R., Pritzel, A., Green, T., Figurnov, M., Ronneberger, O., Tunyasuvunakool, K., Bates, R., Žídek, A., Potapenko, A., et al. (2021). Highly accurate protein structure prediction with AlphaFold. Nature 596, 583–589. 10.1038/s41586-021-03819-2.

53. Kumar, B., Hawkins, G.M., Kicmal, T., Qing, E., Timm, E., and Gallagher, T. (2021). Assembly and Entry of Severe Acute Respiratory Syndrome Coronavirus 2 (SARS-CoV2): Evaluation Using Virus-Like Particles. Cells 10, 853. 10.3390/cells10040853.

54. Sasaki, M., Anindita, P.D., Phongphaew, W., Carr, M., Kobayashi, S., Orba, Y., and Sawa, H. (2018). Development of a rapid and quantitative method for the analysis of viral entry and release using a NanoLuc luciferase complementation assay. Virus Res. 243, 69–74. 10.1016/j.virusres.2017.10.015.

55. Vennema, H., Godeke, G.J., Rossen, J.W., Voorhout, W.F., Horzinek, M.C., Opstelten, D.J., and Rottier, P.J. (1996). Nucleocapsid-independent assembly of coronavirus-like particles by co-expression of viral envelope protein genes. EMBO J. 15, 2020–2028. 10.1002/j.1460-2075.1996.tb00553.x.

56. Baudoux, P., Carrat, C., Besnardeau, L., Charley, B., and Laude, H. (1998). Coronavirus Pseudoparticles Formed with Recombinant M and E Proteins Induce Alpha Interferon Synthesis by Leukocytes. J. Virol. 72, 8636–8643. 10.1128/jvi.72.11.8636-8643.1998.

57. Swann, H., Sharma, A., Preece, B., Peterson, A., Eldredge, C., Belnap, D.M., Vershinin, M., and Saffarian, S. (2020). Minimal system for assembly of SARS-CoV-2 virus like particles. Sci. Rep. 10, 21877. 10.1038/s41598-020-78656-w.

58. García-Murria, M.J., Expósito-Domínguez, N., Duart, G., Mingarro, I., and Martinez-Gil, L. (2019). A Bimolecular Multicellular Complementation System for the Detection of Syncytium Formation: A New Methodology for the Identification of Nipah Virus Entry Inhibitors. Viruses 11, 229. 10.3390/v11030229.

59. Ortiz-Mateu, J., Belda, D., Aviles-Alia, A.I., Alonso-Romero, J., Garcia-Murria, M.J., Mingarro, I., Geller, R., and Martinez-Gil, L. (2025). The sequence and structural integrity of the SARS-CoV-2 Spike protein transmembrane domain is crucial for viral entry. 10.1101/2024.11.19.624333.

60. Xiang, Q., Wu, J., Zhou, Y., Li, L., Tian, M., Li, G., Zhang, Z., and Fu, Y. (2024). SARS-CoV-2 Membrane protein regulates the function of Spike by inhibiting its plasma membrane localization and enzymatic activity of Furin. Microbiol. Res. 282, 127659. 10.1016/j.micres.2024.127659.

61. Perrier, A., Bonnin, A., Desmarets, L., Danneels, A., Goffard, A., Rouillé, Y., Dubuisson, J., and Belouzard, S. (2019). The C-terminal domain of the MERS coronavirus M protein contains a trans-Golgi network localization signal. J. Biol. Chem. 294, 14406–14421. 10.1074/jbc.RA119.008964.

62. Tseng, Y.-T., Wang, S.-M., Huang, K.-J., Lee, A.I.-R., Chiang, C.-C., and Wang, C.-T. (2010). Self-assembly of Severe Acute Respiratory Syndrome Coronavirus Membrane Protein*. J. Biol. Chem. 285, 12862–12872. 10.1074/jbc.M109.030270.

63. Ujike, M., and Taguchi, F. (2015). Incorporation of Spike and Membrane Glycoproteins into Coronavirus Virions. Viruses 7, 1700–1725. 10.3390/v7041700.

64. McDowell, M.A., Heimes, M., Enkavi, G., Farkas, Á., Saar, D., Wild, K., Schwappach, B., Vattulainen, I., and Sinning, I. (2023). The GET insertase exhibits conformational plasticity and induces membrane thinning. Nat. Commun. 14, 7355. 10.1038/s41467-023-42867-2.

65. Bai, L., You, Q., Feng, X., Kovach, A., and Li, H. (2020). Structure of the ER membrane complex, a transmembrane-domain insertase. Nature 584, 475–478. 10.1038/s41586-020-2389-3.

66. Yang, T.-J., Mukherjee, S., Langer, J.D., Hummer, G., and McDowell, M.A. (2025). SND3 is the membrane insertase within a fungal multipass translocon. 10.1101/2025.07.08.663624.

67. Ramírez, A.S., Kowal, J., and Locher, K.P. (2019). Cryo–electron microscopy structures of human oligosaccharyltransferase complexes OST-A and OST-B. Science 366, 1372–1375. 10.1126/science.aaz3505.

68. Coelho, J.P.L., Stahl, M., Bloemeke, N., Meighen-Berger, K., Alvira, C.P., Zhang, Z.-R., Sieber, S.A., and Feige, M.J. (2019). A network of chaperones prevents and detects failures in membrane protein lipid bilayer integration. Nat. Commun. 10, 672. 10.1038/s41467-019-08632-0.

69. Hegde, R.S., and Keenan, R.J. (2022). The mechanisms of integral membrane protein biogenesis. Nat. Rev. Mol. Cell Biol. 23, 107–124. 10.1038/s41580-021-00413-2.

70. Hegde, R.S., and Keenan, R.J. (2024). A unifying model for membrane protein biogenesis. Nat. Struct. Mol. Biol. 31, 1009–1017. 10.1038/s41594-024-01296-5.

71. Mermans, D., Nicolaus, F., Fleisch, K., and von Heijne, G. (2022). Cotranslational folding and assembly of the dimeric Escherichia coli inner membrane protein EmrE. Proc. Natl. Acad. Sci. 119, e2205810119. 10.1073/pnas.2205810119.

72. Du, K., and Lukacs, G.L. (2009). Cooperative Assembly and Misfolding of CFTR Domains In Vivo. Mol. Biol. Cell 20, 1903–1915. 10.1091/mbc.e08-09-0950.

73. van der Sluijs, P., Hoelen, H., Schmidt, A., and Braakman, I. (2024). The Folding Pathway of ABC Transporter CFTR: Effective and Robust. J. Mol. Biol. 436, 168591. 10.1016/j.jmb.2024.168591.

74. Duran-Romaña, R., Houben, B., Migens, P.F., Zhang, Y., Rousseau, F., and Schymkowitz, J. (2025). Native Fold Delay and its implications for co-translational chaperone binding and protein aggregation. Nat. Commun. 16, 1673. 10.1038/s41467-025-57033-z.

75. Aviner, R., and Frydman, J. (2020). Proteostasis in Viral Infection: Unfolding the Complex Virus–Chaperone Interplay. 10.1101/cshperspect.a034090.

76. Wang, N., Seko, A., Takeda, Y., and Ito, Y. (2020). Glycan dependent refolding activity of ER glucosyltransferase (UGGT). Biochim. Biophys. Acta BBA - Gen. Subj. 1864, 129709. 10.1016/j.bbagen.2020.129709.

77. Arnold, S.M., Fessler, L.I., Fessler, J.H., and Kaufman, R.J. (2000). Two Homologues Encoding Human UDP-Glucose:Glycoprotein Glucosyltransferase Differ in mRNA Expression and Enzymatic Activity. Biochemistry 39, 2149–2163. 10.1021/bi9916473.

78. Mann, M.K., Yin, Y., Marsili, S., Xie, J., Doijen, J., Miller, R., Piassek, M., van den Broeck, N., Kariuki, C.K., de Gruyter, H.L.M., et al. (2025). Structural insights into MERS and SARS coronavirus membrane proteins. Commun. Biol. 8, 1651. 10.1038/s42003-025-09042-3.

79. Heaton, N.S., Moshkina, N., Fenouil, R., Gardner, T.J., Aguirre, S., Shah, P.S., Zhao, N., Manganaro, L., Hultquist, J.F., Noel, J., et al. (2016). Targeting Viral Proteostasis Limits Influenza Virus, HIV, and Dengue Virus Infection. Immunity.

80. Pohl, M.O., Martin-Sancho, L., Ratnayake, R., White, K.M., Riva, L., Chen, Q.-Y., Lieber, G., Busnadiego, I., Yin, X., Lin, S., et al. (2022). Sec61 Inhibitor Apratoxin S4 Potently Inhibits SARS-CoV-2 and Exhibits Broad-Spectrum Antiviral Activity. ACS Infect. Dis. 8, 1265– 1279. 10.1021/acsinfecdis.2c00008.

81. Siu, Y.L., Teoh, K.T., Lo, J., Chan, C.M., Kien, F., Escriou, N., Tsao, S.W., Nicholls, J.M., Altmeyer, R., Peiris, J.S.M., et al. (2008). The M, E, and N structural proteins of the severe acute respiratory syndrome coronavirus are required for efficient assembly, trafficking, and release of virus-like particles. J. Virol. 82, 11318–11330. 10.1128/JVI.01052-08.

82. Julius, A., Laur, L., Schanzenbach, C., and Langosch, D. (2017). BLaTM 2.0, a Genetic Tool Revealing Preferred Antiparallel Interaction of Transmembrane Helix 4 of the Dual-Topology Protein EmrE. J. Mol. Biol. 429, 1630–1637. 10.1016/j.jmb.2017.04.003.

83. Lemmon, M.A., Flanagan, J.M., Hunt, J.F., Adair, B.D., Bormann, B.J., Dempsey, C.E., and Engelman, D.M. (1992). Glycophorin A dimerization is driven by specific interactions between transmembrane alphahelices. J. Biol. Chem. 267, 7683–7689. 10.1016/S0021-9258(18)42569-0.

84. MacKenzie, K.R., Prestegard, J.H., and Engelman, D.M. (1997). A Transmembrane Helix Dimer: Structure and Implications. Science 276, 131–133. 10.1126/science.276.5309.131.

85. Orzáez, M., Pérez-Payá, E., and Mingarro, I. (2000). Influence of the C-terminus of the glycophorin A transmembrane fragment on the dimerization process. Protein Sci. 9, 1246–1253. 10.1110/ps.9.6.1246.

86. Srinivasan, S., Deng, W., and Li, R. (2011). L-selectin transmembrane and cytoplasmic domains are monomeric in membranes. Biochim. Biophys. Acta BBA - Biomembr. 1808, 1709–1715. 10.1016/j.bbamem.2011.02.006.

87. Andreu-Fernández, V., Sancho, M., Genovés, A., Lucendo, E., Todt, F., Lauterwasser, J., Funk, K., Jahreis, G., Pérez-Payá, E., Mingarro, I., et al. (2017). Bax transmembrane domain interacts with prosurvival Bcl-2 proteins in biological membranes. Proc. Natl. Acad. Sci. 114, 310– 315. 10.1073/pnas.1612322114.

88. Abramson, J., Adler, J., Dunger, J., Evans, R., Green, T., Pritzel, A., Ronneberger, O., Willmore, L., Ballard, A.J., Bambrick, J., et al. (2024). Accurate structure prediction of biomolecular interactions with Al-phaFold 3. Nature 630, 493–500. 10.1038/s41586-024-07487-w.

89. Jo, S., Kim, T., Iyer, V.G., and Im, W. (2008). CHARMM-GUI: A web-based graphical user interface for CHARMM. J. Comput. Chem. 29, 1859–1865. 10.1002/jcc.20945.

90. Phillips, J.C., Hardy, D.J., Maia, J.D.C., Stone, J.E., Ribeiro, J.V., Bernardi, R.C., Buch, R., Fiorin, G., Hénin, J., Jiang, W., et al. (2020). Scalable molecular dynamics on CPU and GPU architectures with NAMD. J. Chem. Phys. 153, 044130. 10.1063/5.0014475.

91. Huang, J., Rauscher, S., Nawrocki, G., Ran, T., Feig, M., de Groot, B.L., Grubmüller, H., and MacKerell, A.D. (2017). CHARMM36m: an improved force field for folded and intrinsically disordered proteins. Nat. Methods 14, 71–73. 10.1038/nmeth.4067.

92. Jorgensen, W.L., Chandrasekhar, J., Madura, J.D., Impey, R.W., and Klein, M.L. (1983). Comparison of simple potential functions for simulating liquid water. J. Chem. Phys. 79, 926–935. 10.1063/1.445869.

93. Darden, T., York, D., and Pedersen, L. (1993). Particle mesh Ewald: An N·log(N) method for Ewald sums in large systems. J. Chem. Phys. 98, 10089–10092. 10.1063/1.464397.

94. Yamamoto, M., Du, Q., Song, J., Wang, H., Watanabe, A., Tanaka, Y., Kawaguchi, Y., Inoue, J., and Matsuda, Z. (2019). Cell–cell and virus– cell fusion assay–based analyses of alanine insertion mutants in the distal α9 portion of the JRFL gp41 subunit from HIV-1. J. Biol. Chem. 294, 5677–5687. 10.1074/jbc.RA118.004579.

95. Hoffmann, M., Kleine-Weber, H., Schroeder, S., Krüger, N., Herrler, T., Erichsen, S., Schiergens, T.S., Herrler, G., Wu, N.-H., Nitsche, A., et al. (2020). SARS-CoV-2 Cell Entry Depends on ACE2 and TMPRSS2 and Is Blocked by a Clinically Proven Protease Inhibitor. Cell 181, 271-280.e8. 10.1016/j.cell.2020.02.052.

96. Branon, T.C., Bosch, J.A., Sanchez, A.D., Udeshi, N.D., Svinkina, T., Carr, S.A., Feldman, J.L., Perrimon, N., and Ting, A.Y. (2018). Efficient proximity labeling in living cells and organisms with TurboID. Nat. Biotechnol. 36, 880–887. 10.1038/nbt.4201.

97. Gingras, A.-C., Abe, K.T., and Raught, B. (2019). Getting to know the neighborhood: using proximity-dependent biotinylation to characterize protein complexes and map organelles. Curr. Opin. Chem. Biol. 48, 44–54. 10.1016/j.cbpa.2018.10.017.

98. Laurent, E.M.N., Sofianatos, Y., Komarova, A., Gimeno, J.-P., Tehrani, P.S., Kim, D.-K., Abdouni, H., Duhamel, M., Cassonnet, P., Knapp, J.J., et al. (2020). Global BioID-based SARS-CoV-2 proteins proximal interactome unveils novel ties between viral polypeptides and host factors involved in multiple COVID19-associated mechanisms. Preprint at bioRxiv, 10.1101/2020.08.28.272955 https://doi.org/10.1101/2020.08.28.272955.

99. Lee, Y.-B., Jung, M., Kim, J., Charles, A., Christ, W., Kang, J., Kang, M.-G., Kwak, C., Klingström, J., Smed-Sörensen, A., et al. (2023). Super-resolution proximity labeling reveals anti-viral protein network and its structural changes against SARS-CoV-2 viral proteins. Cell Rep. 42. 10.1016/j.celrep.2023.112835.

100. Zhang, Y., Shang, L., Zhang, J., Liu, Y., Jin, C., Zhao, Y., Lei, X., Wang, W., Xiao, X., Zhang, X., et al. (2022). An antibody-based proximity labeling map reveals mechanisms of SARS-CoV-2 inhibition of antiviral immunity. Cell Chem. Biol. 29, 5-18.e6. 10.1016/j.chembiol.2021.10.008.

101. Kim, K., Park, I., Kim, J., Kang, M.-G., Choi, W.G., Shin, H., Kim, J.-S., Rhee, H.-W., and Suh, J.M. (2021). Dynamic tracking and identification of tissue-specific secretory proteins in the circulation of live mice. Nat. Commun. 12, 5204. 10.1038/s41467-021-25546-y.

102. Cox, J., and Mann, M. (2008). MaxQuant enables high peptide identification rates, individualized p.p.b.-range mass accuracies and proteome-wide protein quantification. Nat. Biotechnol. 26, 1367–1372. 10.1038/nbt.1511.

103. Cox, J., Neuhauser, N., Michalski, A., Scheltema, R.A., Olsen, J.V., and Mann, M. (2011). Andromeda: A Peptide Search Engine Integrated into the MaxQuant Environment. J. Proteome Res. 10, 1794–1805. 10.1021/pr101065j.

104. Shannon, P., Markiel, A., Ozier, O., Baliga, N.S., Wang, J.T., Ramage, D., Amin, N., Schwikowski, B., and Ideker, T. (2003). Cytoscape: A Software Environment for Integrated Models of Biomolecular Interaction Networks. 10.1101/gr.1239303.

105. Morris, J.H., Apeltsin, L., Newman, A.M., Baumbach, J., Wittkop, T., Su, G., Bader, G.D., and Ferrin, T.E. (2011). clusterMaker: a multi-algorithm clustering plugin for Cytoscape. BMC Bioinformatics 12, 436. 10.1186/1471-2105-12-436.

106. Doncheva, N.T., Morris, J.H., Gorodkin, J., and Jensen, L.J. (2019). Cytoscape StringApp: Network Analysis and Visualization of Proteomics Data. J. Proteome Res. 18, 623–632. 10.1021/acs.jproteome.8b00702.

107. Madeira, F., Park, Y. mi, Lee, J., Buso, N., Gur, T., Madhusoodanan, N., Basutkar, P., Tivey, A.R.N., Potter, S.C., Finn, R.D., et al. (2019). The EMBL-EBI search and sequence analysis tools APIs in 2019. Nucleic Acids Res. 47, W636–W641. 10.1093/nar/gkz268.

108. Waterhouse, A.M., Procter, J.B., Martin, D.M.A., Clamp, M., and Barton, G.J. (2009). Jalview Version 2—a multiple sequence alignment editor and analysis workbench. Bioinformatics 25, 1189–1191. 10.1093/bioinformatics/btp033.

