## Supplementary Figures for "SARS-CoV-2 membrane protein biogenesis"

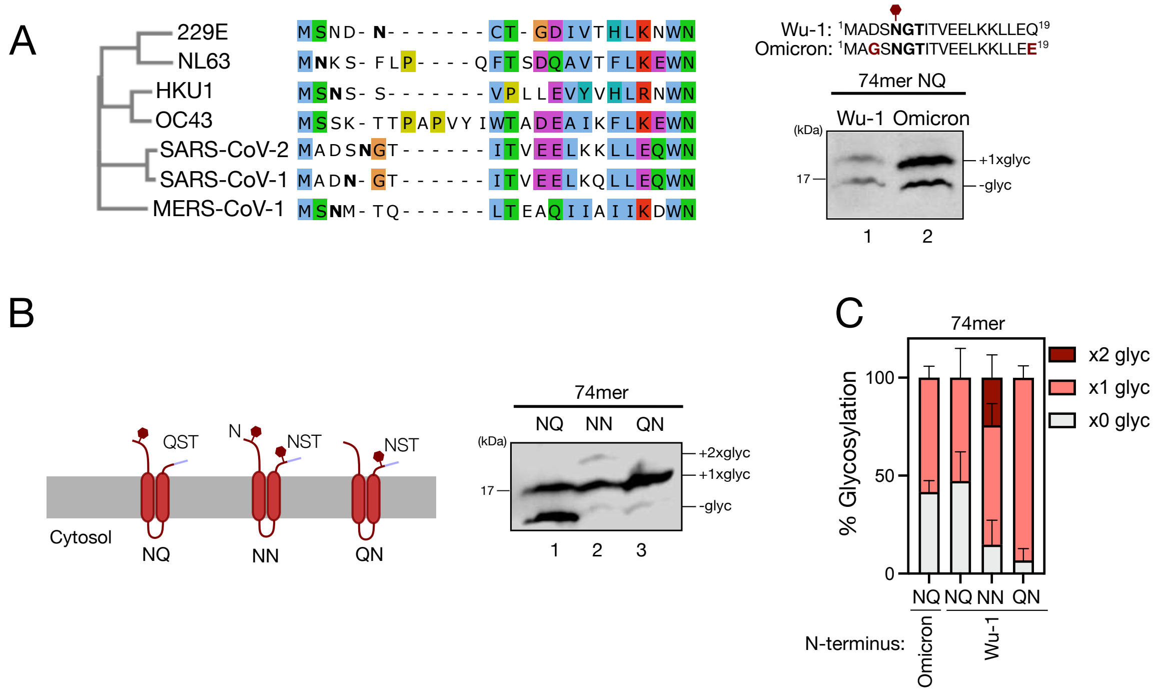


**Figure S1. SARS-CoV-2 M protein TMD2 insertion.** (A) Sequence alignment of the N-terminal domain of the M protein across human coronaviruses. Sequences are colored using the Clustal X color scheme, where white residues are non-conserved. Incorporating the Omicron-specific D3G and Q19E mutations into the N-terminus did not significantly affect glycosylation efficiency. (B) Glycosylation efficiency of the natural NGT site. To explore the glycosylation efficiency of the native M site, and whether it depends on OST activity or its sequence, we designed different glycosylation sites for the M protein 74mer truncate. The QST (NQ) and NST (NN) constructs including the native site were included, along with a construct preserving the NST C-terminal site but with a mutated native site (N5Q, QN). We confirmed the relatively low glycosylation efficiency of the native site (NQ, 53%) compared to the designed site at the C-terminus (QN, 97%). (C) Quantification of glycosylated and not-glycosylated bands (mean ± SD; N = 5–9).

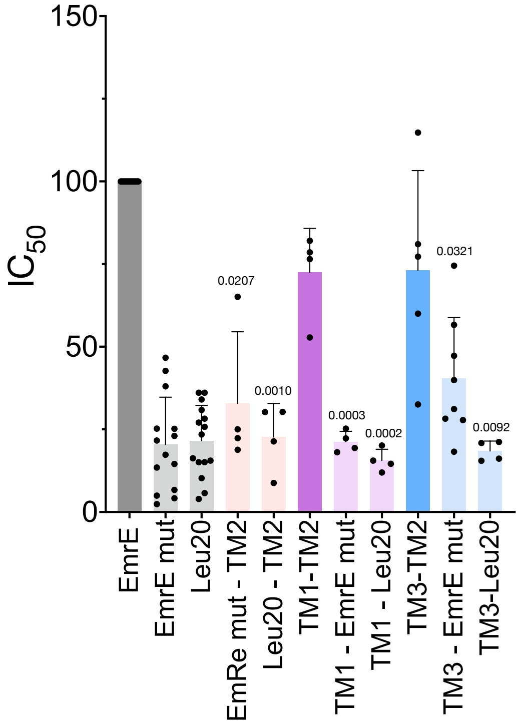


**Figure S2. BLaTM control TMD–TMD interactions.** LD50 values for heterotypic interactions between M protein TMDs and various negative controls were assessed using the BlaTM assay. Both EmrE mutant version and Leu20 were tested for interactions with M TMDs using BLaTM 2.0. Interactions between TMD2 and TMD1 (purple) or TMD3 (blue) were used as positive controls. The system does not support testing antiparallel TMD–TMD interactions; thus, negative controls for TMD2 interactions were compared against the EmrE positive control. All values are normalized to the respective positive controls (dark colors; set to 100%). Data are presented as mean ± SD. Each solid dot represents an individual experiment (N = 4–15). Above bars: p values below the significance threshold (<0.05) calculated using one-sample t-tests against a reference value of "100".

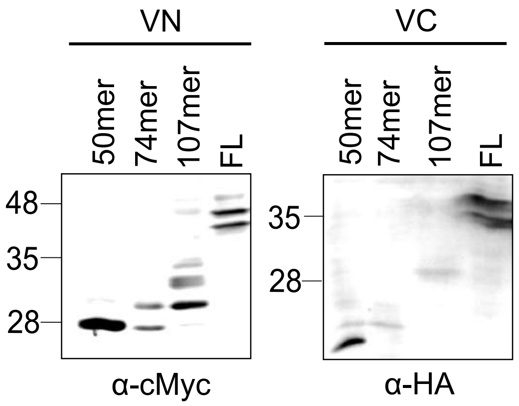


**Figure S3. Transient expression of M-split VFP chimeras in human cells** Western blot analysis of the expression levels of M-VN and M-VC truncated variants in HEK293T cells.

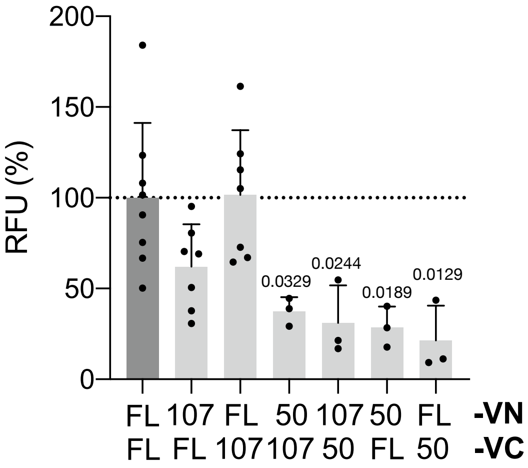


**Figure S4. Bimolecular fluorescence complementation (BiFC) signal for heterotypic interactions between M-protein truncations and the full-length construct.** Relative fluorescence units (RFU) are shown for each interaction pair. All values are normalized to the mean RFU of the full-length M homo-interaction (FL, dark gray; set to 100%). Data are presented as mean ± SD. Each solid dot represents an individual experiment (N = 3–8). Above bars: p values below the significance threshold (<0.05), calculated using one-sample t-tests against a reference value of 100.

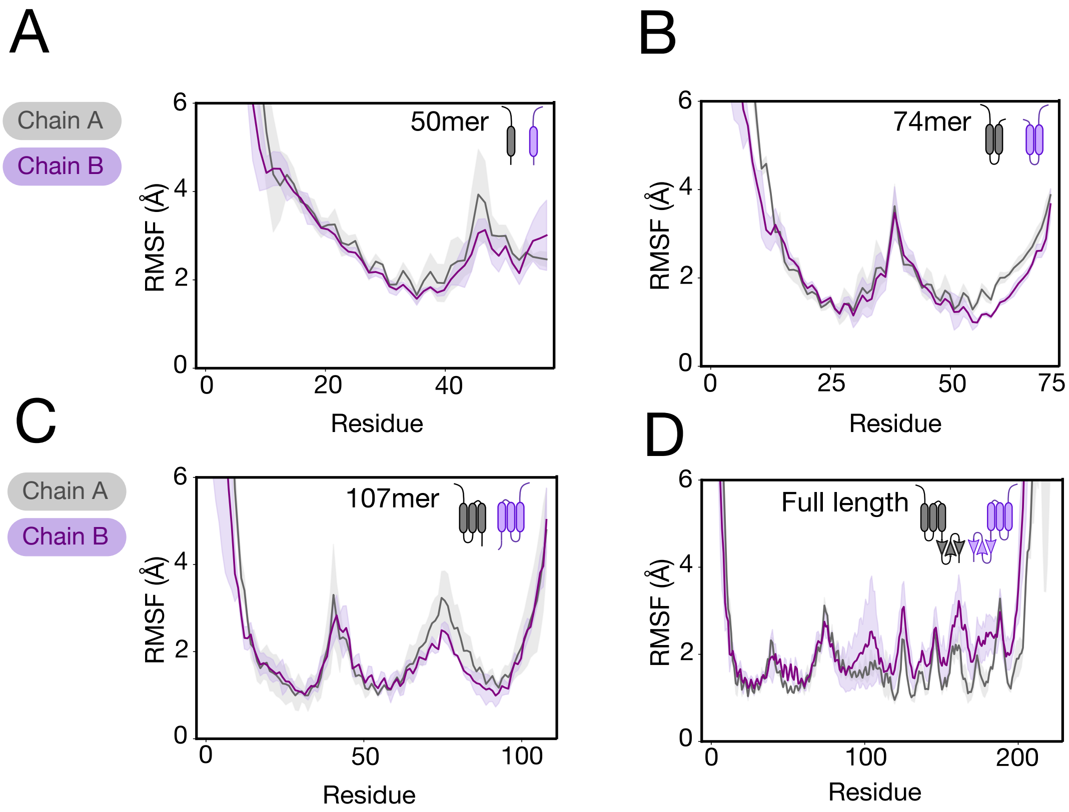


**Figure S5. Analysis of root mean square fluctuation (RMSF) of atomic positions for dimer constructs from MD simulations in the ER membrane.** (A) 50mer. (B) 74mer. (C) 107mer. (D) Full-length. Solid lines represent the mean RMSF. Shaded areas indicate the standard deviation across three

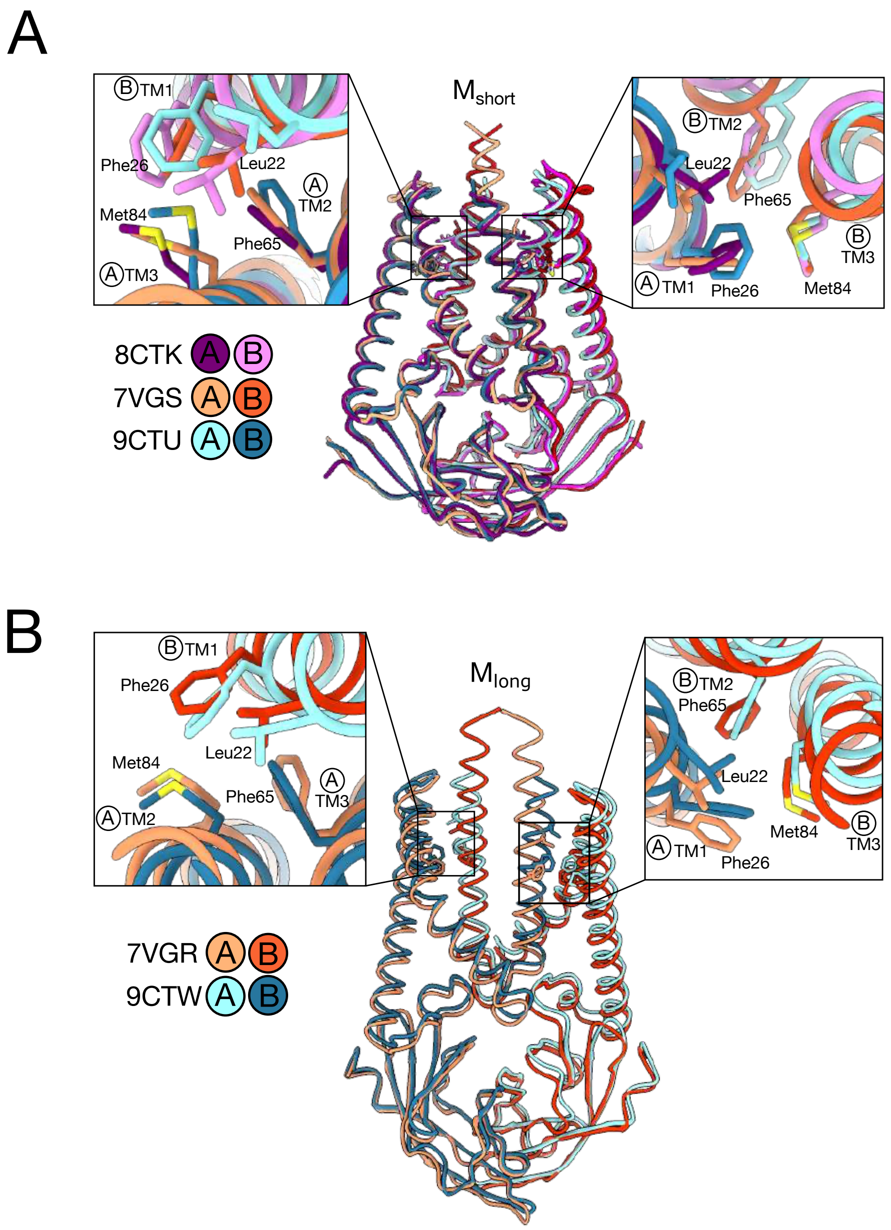


**Figure S6. Structural analysis of a transmembrane hydrophobic cluster in SARS-CoV-2 M cryo-EM structures.** (A) Overlay of available cryo-EM structures of the M protein "short" isoform from the PDB (IDs 8CTK, 7VGS, and 9CTU). A cluster of four hydrophobic residues is highlighted and enlarged. (B) Same as in (A) for the "long" isoform of the M protein (PDB IDs codes 7VGR and 9CTW).

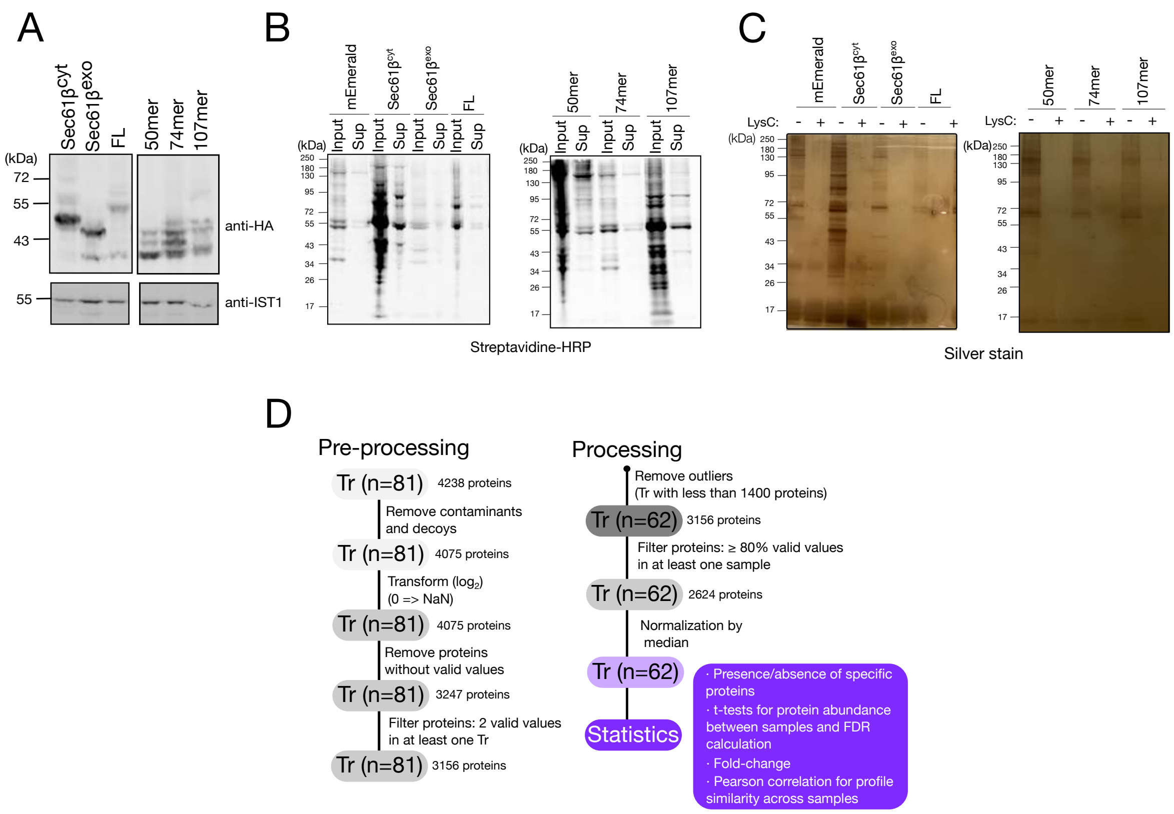


**Figure S7. Quality control and data processing in the proximity labeling workflow.** (A) Expression of HA-TurboID constructs in HEK293T cells. Western blot analysis of HA-TurboID fusion protein expression, with IST-1 used as a loading control. (B) Quality control of TurboID biotinylation and neutravidin pull-down. Samples from the input and supernatant following neutravidin bead incubation were resolved by SDS-PAGE, and probed with Streptavidin-HRP to assess biotinylation efficiency and pull-down performance. (C) Evaluation of protein recovery before and after LysC cleavage. Proteins captured on neutravidin beads were eluted by LysC digestion or boiling in Laemmli buffer with β-mercaptoethanol. Samples were resolved by SDS-PAGE, and visualized by silver staining. (D) Schematic view of the processing pipeline used to select bona fide Turbo-ID tagged proteins, and to identify the differential interactome of each chimeric protein. Numbers in parenthesis indicate the total number of LC-MS/MS samples analyzed in each step.

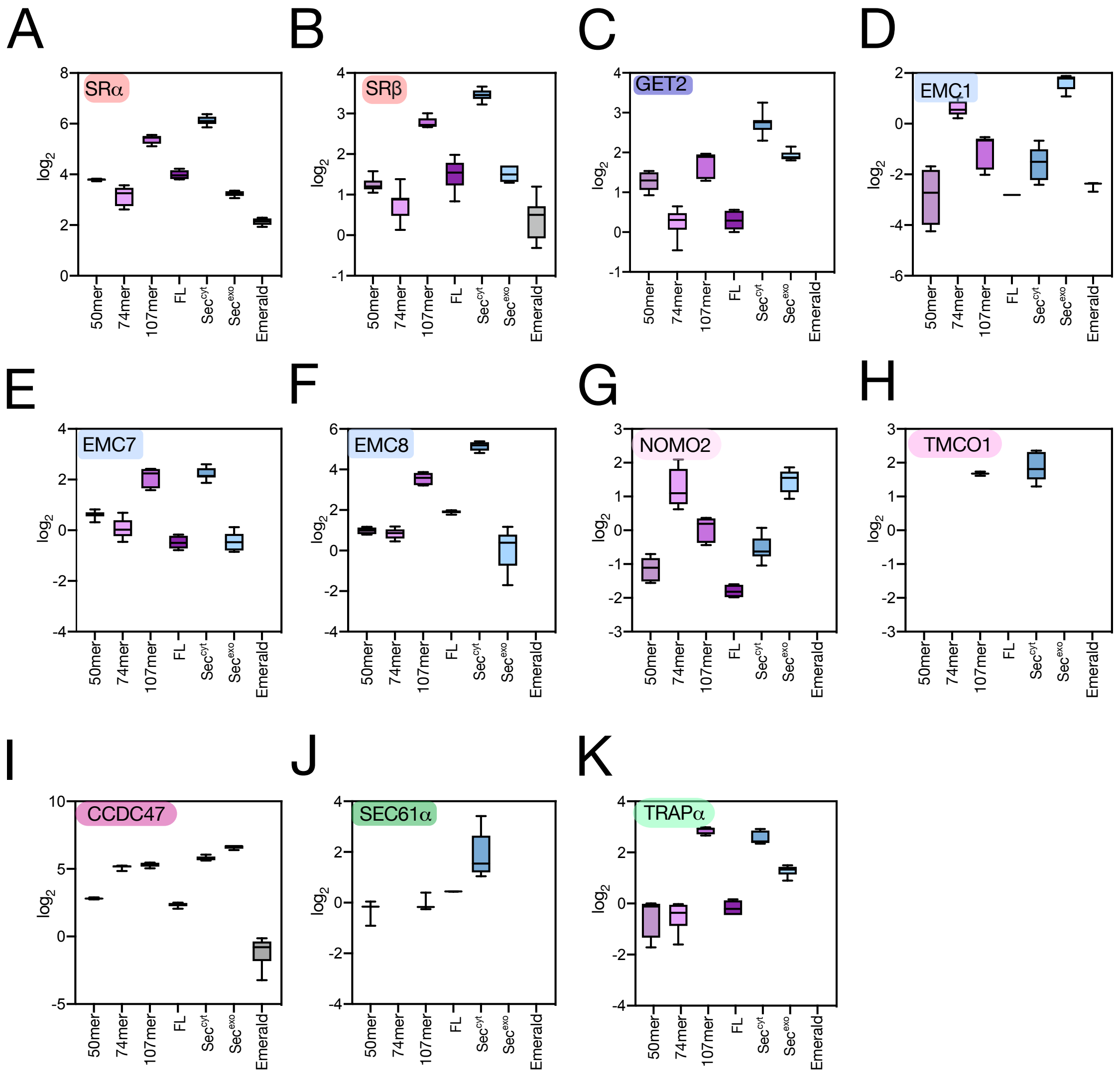


**Figure S8. LFQ abundance values for ER targeting and insertion human cofactors .** (A) Box plot showing the detailed abundance of SRα protein detected in our MS data. Protein abundance for each condition is represented as the median normalized (represented as the '0' value) in log2 scale. Whiskers show minimum to maximum values. Boxes extend from the 25th to the 75th percentiles. Middle line depicts the median LFQ value for each specific sample. (B) Same as in (A) for SRβ protein. (C) Same as in (A) for GET2 protein. (D) Same as in (A) for EMC1 protein. (E) Same as in (A) for EMC7 protein. (F) Same as in (A) for EMC8 protein. (G) Same as in (A) for NOMO2 protein. (H) Same as in (A) for TMCO1 protein. (I) Same as in (A) for CCDC47 protein. (J) Same as in (A) for SEC61α protein. (K) Same as in (A) for TRAPα protein.

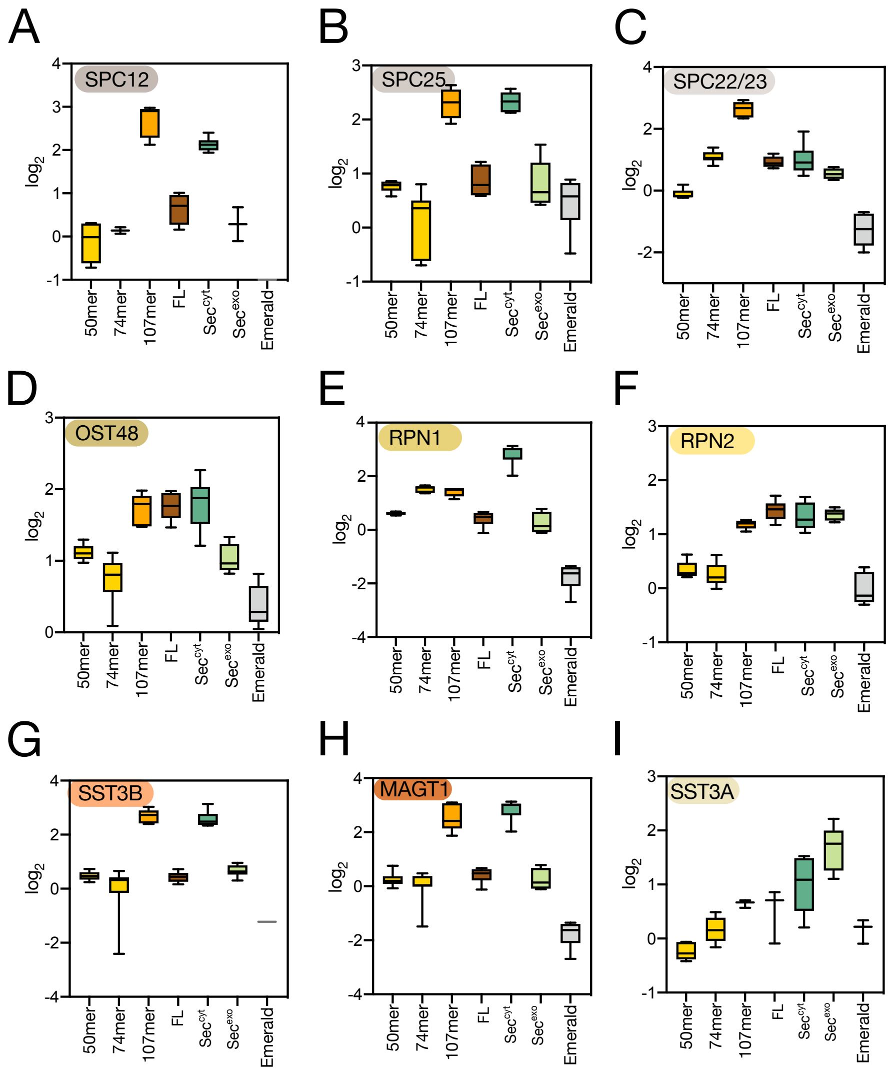


**Figure S9. LFQ abundance values for other ER protein biogenesis human cofactors**. (A) Box plot showing the detailed abundance of SPC12 proteins detected in our MS data. Protein abundance for each condition is represented as the median normalized in log2 scale. Whiskers show minimum to maximum values. Boxes extend from the 25th to the 75th percentiles. Middle line depicts the median. (B) Same as in (A) for SPC25 protein. (C) Same as in (A) for SPC22/23 protein. (D) Same as in (A) for OST48 protein. (E) Same as in (A) for RPN1 protein. (F) Same as in (A) for RPN2 protein. (G) Same as in (A) for SST3B protein. (H) Same as in (A) for MAGT1 protein. (I) Same as in (A) for SST3A protein.

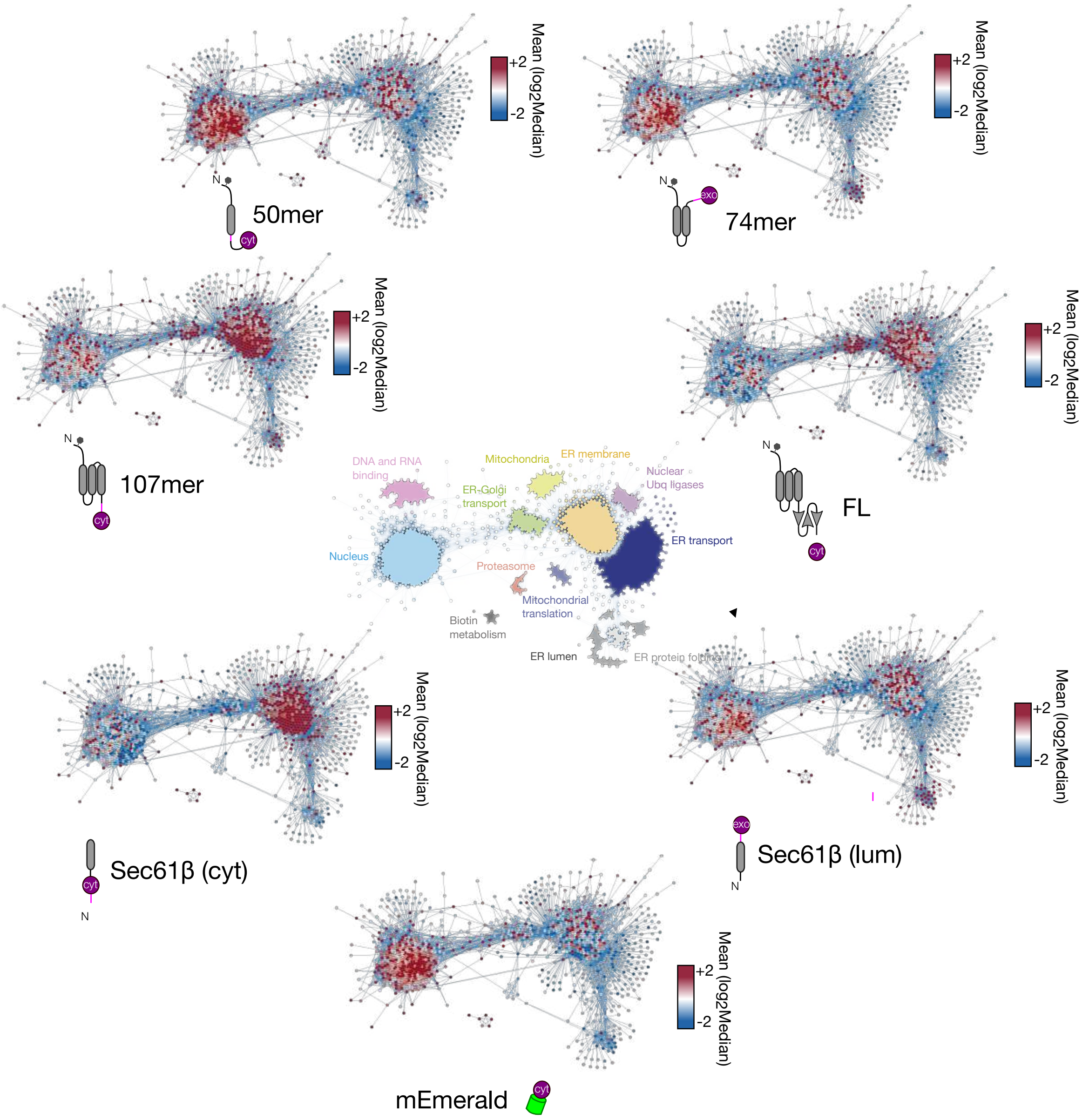


**Figure S10. Protein abundance within correlation networks.** The proteins in the correlation network shown in Figure 6, colored according to the log2 of the normalized abundance of the samples indicated in the labels. The most abundant proteins are colored red. The least abundant are colored blue.
